# B cell receptor induced IL-10 production from neonatal CD19+CD43- cells depends on STAT5 mediated IL-6 secretion

**DOI:** 10.1101/2022.09.18.508441

**Authors:** Jiro Sakai, Jiyeon Yang, Chao-Kai Chou, Wells W. Wu, Mustafa Akkoyunlu

## Abstract

Newborns are unable to reach the adult-level humoral immune response partly due to the potent immunoregulatory role of IL-10. Increased IL-10 production by neonatal B cells has been attributed to the larger population of IL-10-producting CD43^+^ B-1 cells in neonates. Here, we show that neonatal CD43^-^ non-B-1 cells also produce substantial amounts of IL-10 following B cell antigen receptor (BCR) activation. In neonatal CD43^-^ non-B-1 cells, BCR engagement activated STAT5 under the control of phosphorylated forms of signaling molecules Syk, Btk, PKC, FAK and Rac1. STAT5 activation led to IL-6 production, which in turn was responsible for IL-10 production in an autocrine/paracrine fashion through the activation of STAT3. In addition to the increased IL-6 production in response to BCR stimulation, elevated expression of IL-6Rα expression in neonatal B cells rendered them highly susceptible to IL-6 mediated STAT3 phosphorylation and IL-10 production. Finally, IL-10 secreted from neonatal CD43^-^ non-B-1 cells was sufficient to inhibit TNF-α secretion by macrophages. Our results unveil a distinct mechanism of IL-6-dependent IL-10 production in BCR-stimulated neonatal CD19^+^CD43^-^ B cells.

## INTRODUCTION

Infants and neonates demonstrate high susceptibility to infection, leading to 40% of the annual death of approximately 7 million children under the age of 5 years worldwide (*1*). Fetomaternal immune tolerance is essential to suppress rejection during pregnancy, and disruption of the fetomaternal tolerance can result in preterm labor (*2*). During the perinatal period, newborns transition from their dependence on maternal immunity to their own immune system (*3*). Rapid exposure to environmental assaults, such as microbes, after birth renders neonates susceptible to infections (*4, 5*). Several types of immune suppressive cells contribute to fetomaternal immune tolerance and to neonatal suboptimal immunity via the potent anti-inflammatory cytokine interleukin (IL)-10 (*5, 6*).

Initially described as a function of activated CD4^+^ T helper (Th) 2 cells to inhibit cytokine production by Th1 cells (*7*), IL-10 has subsequently been shown to be produced by different cell types including dendritic cells, macrophages, neutrophils, mast cells, natural killer cells, T cells, and B cells (*8, 9*). The main activity of IL-10 is the inhibition of inflammatory responses (e.g., pro-inflammatory cytokine and chemokine synthesis, nitric oxide production, and antigen presentation) by both the innate and the adaptive immune cells (*8, 9*). IL-10 expression is controlled by various transcription factors including nuclear factor-κB (NF-κB), signal transducer and activator of transcription 3 (STAT3), and GATA binding protein 3 (GATA3) depending on the upstream signaling pathways and cell types (*10, 11*). In B cells, IL-10 production has been shown to be mediated by B cell receptor (BCR) engagement (*12*), CD40 ligand (*13*), TLR agonists (*14*), IL-1β and IL-6 (*15*).

B cells can be divided into 2 subsets: CD43^+^ B-1 cells and CD43^-^ non B-1 cells which include follicular B-2 cells, marginal zone B cells and CD5^+^CD1d^hi^ regulatory B cells (Bregs) (*16*). B-1 cells are further subdivided into innate-like B-1a (CD5^+^) cells and B-1b (CD5^-^) cells. After emerging from the liver in the very early stage of life, B-1 cells have been suggested to be maintained by a “self-renewal” process, whereas B-2 cells are continuously produced from progenitors in the bone marrow after birth (*16, 17*). Accordingly, B-1 cells, which dominate the B cell population in neonatal spleen (30% of IgM^+^ splenic B cells), are gradually exceeded by B-2 cells with age and account for approximately 2% in adult splenic B cells (*18, 19*). Among the B cells, B-1 cells are the main producers of IL-10 (*20, 21*) and since B-1 cells account for 1/3 of neonatal mouse spleen, they are considered as the main source of IL-10 in neonatal spleen. Both mouse neonatal splenic and human cord blood Breg cells are shown to manifest suppressive activity (*21–23*).

Here, we demonstrated that there is a sizable splenic CD19^+^CD43^-^ non-B-1 cell population with potent IL-10 producing capacity in response to BCR stimulation. Upon BCR stimulation, neonatal CD19^+^CD43^-^ cells, but not their adult counterparts, produced IL-6 downstream of activated STAT5. In an autocrine and paracrine fashion, the secreted IL-6 triggered the induction of IL-10 production from CD19^+^CD43^-^ cells. Further, neonatal CD19^+^CD43^-^ cells suppressed inflammatory cytokine production by macrophages in an IL-10-dependent manner.

## RESULTS

### Neonatal spleen contains a CD43^-^ cell population with substantial IL-10 production in response to BCR stimulation

The IL-10-producing splenic B cell population includes several different subsets characterized by surface markers (*24*). Among these subsets, the CD5^+^ B-1 B cells comprise a small portion of adult spleen, but represent a higher percentage of neonatal splenic B cells (*18*). Accordingly, the higher representation of IL-10^+^ B cells in neonatal spleen following a variety of stimuli is thought to be due to higher percentage of CD5^+^ B-1 cells in neonatal spleen (*25, 26*). B-1 cells are identified by CD43 expression, and CD5 is used to distinguish between B-1a (CD43^+^CD5^+^) and B-1b (CD43^+^CD5^-^) (*27, 28*). In this study, we sought to characterize the generation of neonatal IL-10-producing B cells following BCR stimulation. First, we purified CD19^+^ cells from splenocytes (fig. S1). Intracellular IL-10 staining confirmed the increased emergence of CD19^+^IL-10^+^ B cells after stimulation of purified neonatal splenic B cells with anti-IgM antibodies compared to adult B cells (Fig. 1A). Also confirming previous reports (*18, 25*), neonatal spleen contained higher percentage of CD43^+^ B-1 cells than adult spleen (fig. S2). Further gating of purified B cells based on CD43 expression indicated that neonatal CD43^+^ cells contained higher percentage of CD19^+^IL-10^+^ B cells than adult cells prior to BCR stimulation and there was a modest increase after BCR stimulation in both the age groups (Fig. 1B). Unlike the CD43^+^ population, BCR stimulation induced a significant increase in IL-10^+^ cells among the neonatal CD43^-^ population, whereas IL-10^+^ cells remained low in adult CD43^-^ population after BCR-stimulation (Fig. 1B). Thus, although neonatal splenic CD19^+^CD43^+^ cells comprise the main IL-10^+^ cells in neonatal spleen as reported previously, there is a substantial increase in IL-10^+^ cell frequency and mean fluorescence intensity (MFI) among the CD19^+^CD43^-^ population following antigen recognition (Fig. 1B and fig. S3). Since the CD19^+^CD43^+^ cells have been well recognized as the IL-10-producing subset in neonatal spleen, we focused on the characterization of this newly defined IL-10-producing CD19^+^CD43^-^ cell population in neonatal mice. We purified CD19^+^CD43^-^ cells from splenocytes (fig. S4) and measured secreted IL-10 in the culture media of purified CD19^+^CD43^-^ cells after stimulating with anti-IgM antibodies. As observed in the flow cytometry analysis of gated CD19^+^CD43^-^ cells (Fig. 1B), purified neonatal CD19^+^CD43^-^ cells also produced higher IL-10 than the adult counterparts (Fig. 1C). The increase in IL-10 production reached statistical significance at 6-hour time point and remained high after peaking at 12 hours. IL-10 production by adult CD19^+^CD43^-^ cells was minimal at all time points. *Il10* mRNA measurement indicated that adult CD19^+^CD43^-^ cells manifested a rapid and sharp increase in *Il10* expression which was quickly shut off within 6 hours post-stimulation (Fig. 1D), and this temporal gene expression appeared to be insufficient for IL-10 protein synthesis (Fig. 1C). An initial sharp increase of *Il10* expression was also observed in neonatal CD19^+^CD43^-^ cells (Fig. 1D). Unlike in adult cells, neonatal *Il10* expression further increased at later time points, facilitating the production of IL-10 (Fig. 1C). These results indicate that, unlike its adult counterparts, neonatal splenic CD19^+^CD43^-^ cells have the propensity to produce IL-10 after BCR cross-linking.

**Figure 1.**
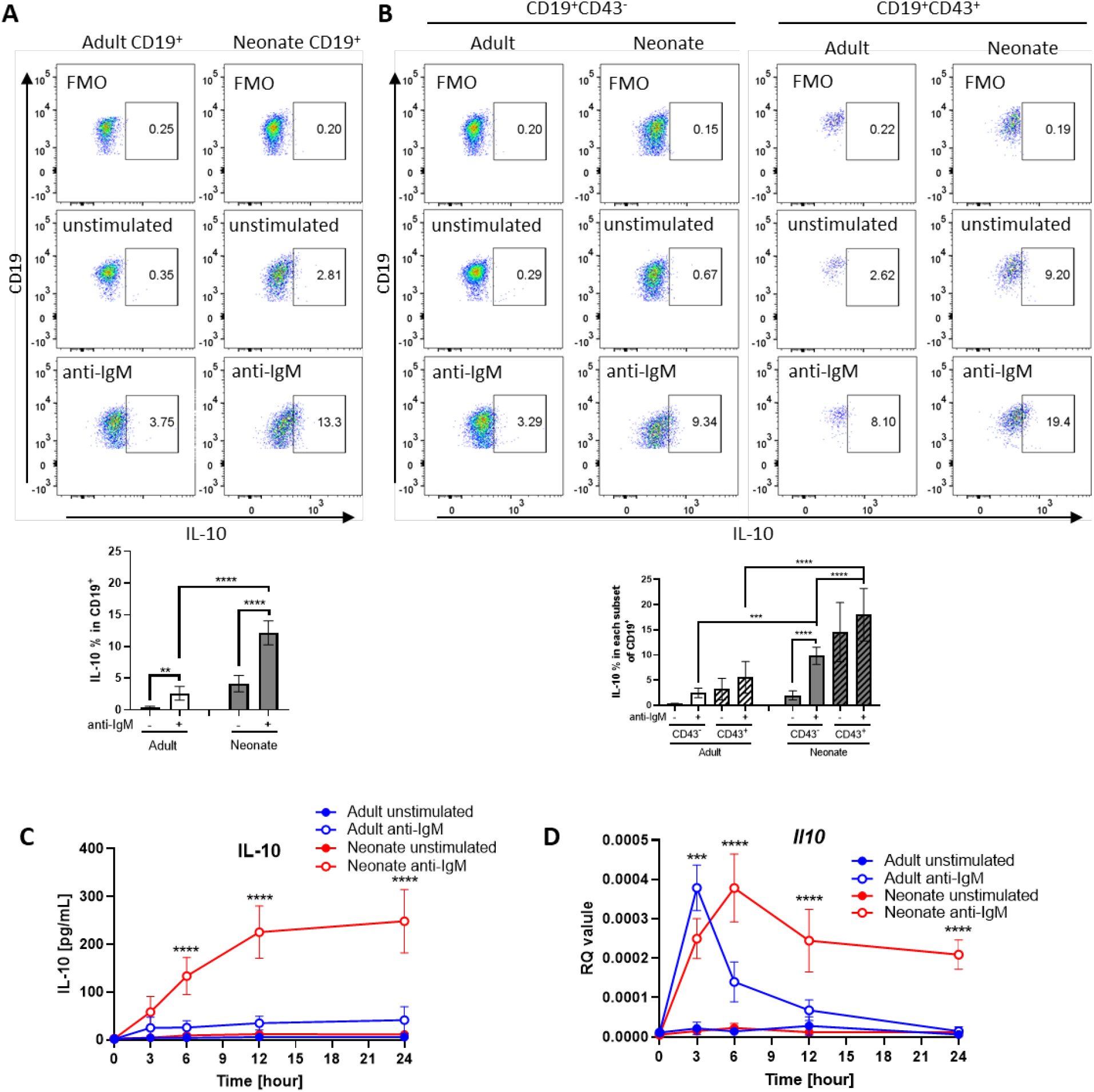
BCR-induced IL-10 production in adult and neonatal splenic B cell subsets. (**A** and **B**) Splenic B cells were isolated from adult and neonatal mice. Isolated cells were incubated in the absence (unstimulated) or presence of 10 μg/mL F(ab’)2 fragments of anti-IgM antibodies for 20 h and intracellular IL-10 production was measured by flow cytometry in gated B cell subsets (*n* = 3). Data are shown as the mean ± s.d. of three independent experiments. *P* values were calculated using one-way ANOVA with a Dunnett’s multiple comparisons test (\*\**P*<0.01, *** *P*<0.001, and **** *P*<0.0001). (**C** and **D**) Isolated adult and neonatal CD19^+^CD43^-^ B cells were incubated at 3 million cells/mL in the absence or presence of 10 μg/mL F(ab’)2 fragments of anti-IgM antibodies for indicated duration and culture supernatant IL-10 levels (C) and mRNA expression (D) were measured in ELISA and q-PCR, respectively. Data are shown as the mean ± s.d. of three independent experiments. *P* values (Adult anti-IgM vs Neonate anti-IgM) were calculated using two-way ANOVA (\*\*\**P*<0.001 and \*\*\*\**P*<0.0001).

### Neonatal BCRs uniquely activate STAT3 and STAT5 following BCR cross-linking

To gain insight into the underlying mechanisms of enhanced IL-10 production in BCR-stimulated neonatal CD19^+^CD43^-^ cells, we performed RNA sequencing (RNA-seq) and Gene Set Enrichment Analysis (GSEA) (*29*) for adult and neonatal B cells following BCR engagement. There were 1,341 increased and 1,905 decreased genes in adult B cells compared to 716 increased and 962 decreased genes in neonatal B cells (fig. S5A). To identify signaling pathways uniquely activated by neonatal BCR, we compared adult and neonatal CD19^+^CD43^-^ cell data using hallmark gene sets from the Molecular Signatures database (MSigDB) C5 gene ontology (GO) collection (*30*). Pathways enriched in neonatal CD19^+^CD43^-^ cells included cytokine receptor signaling pathways leading to STAT protein activation (*31*), whereas adult CD19^+^CD43^-^ cells were enriched for genes involved in biological processes as well as the signaling pathways controlled by mitogen-activated protein kinases (MAPKs) and Akt (*32–37*) (fig. S5B). We also conducted GSEA using the MSigDB C3 transcription factor targets (TFT) hallmark gene collection to identify transcription factors uniquely activated in neonatal B cells. This analysis revealed that target gene sets for STAT3, STAT5A and STAT5B were highly enriched in neonatal B cells, suggesting that these STAT proteins were activated in neonatal CD19^+^CD43^-^ cells after BCR engagement (Fig. 2A and fig. S5C). Supporting the findings in C5 gene ontology analysis, target gene sets such as MYC, ELK1 and AP-1 for MAPK and Akt signaling pathway-activated transcription factors (*38, 39*) were not enriched in neonatal CD19^+^CD43^-^ cells compared to adult cells (Fig. 2A and fig. S5C). To verify the signaling pathways identified by the GSEA analysis of the RNA-seq data, we subjected the BCR-stimulated neonatal and adult CD19^+^CD43^-^ cells to Western blot analysis. We found that STAT3 and STAT5 were highly phosphorylated in neonatal B cells although their activation kinetics was different (Fig. 2, B and C). STAT5 phosphorylation was detected as early as 5 minutes and peaked at 15 minutes after BCR crosslinking (Fig. 2B), whereas STAT3 phosphorylation peaked at around 4 hours post-stimulation (Fig. 2C). Increases in STAT5 and STAT3 phosphorylations were also observed by flow cytometry at 15 minutes and 4 hours post-stimulation, respectively (fig. S5, D and E). There was no significant difference in the enrichment of target gene sets for STAT1 between adult and neonatal B cells (Fig. 2A and fig. S5C). In addition, STAT1 phosphorylation was not observed after BCR cross-linking in either adult or neonatal B cells (Fig. 2, B and C), suggesting no role for STAT1 in BCR-induced IL-10 production. Confirming the GSEA analysis data, Akt, p38, JNK and ERK were phosphorylated in adult B cells, but not in neonatal cells (fig. S6). Collectively, RNA-seq and Western blot analyses revealed important differences between neonatal and adult BCR signaling pathways.

**Figure 2.**
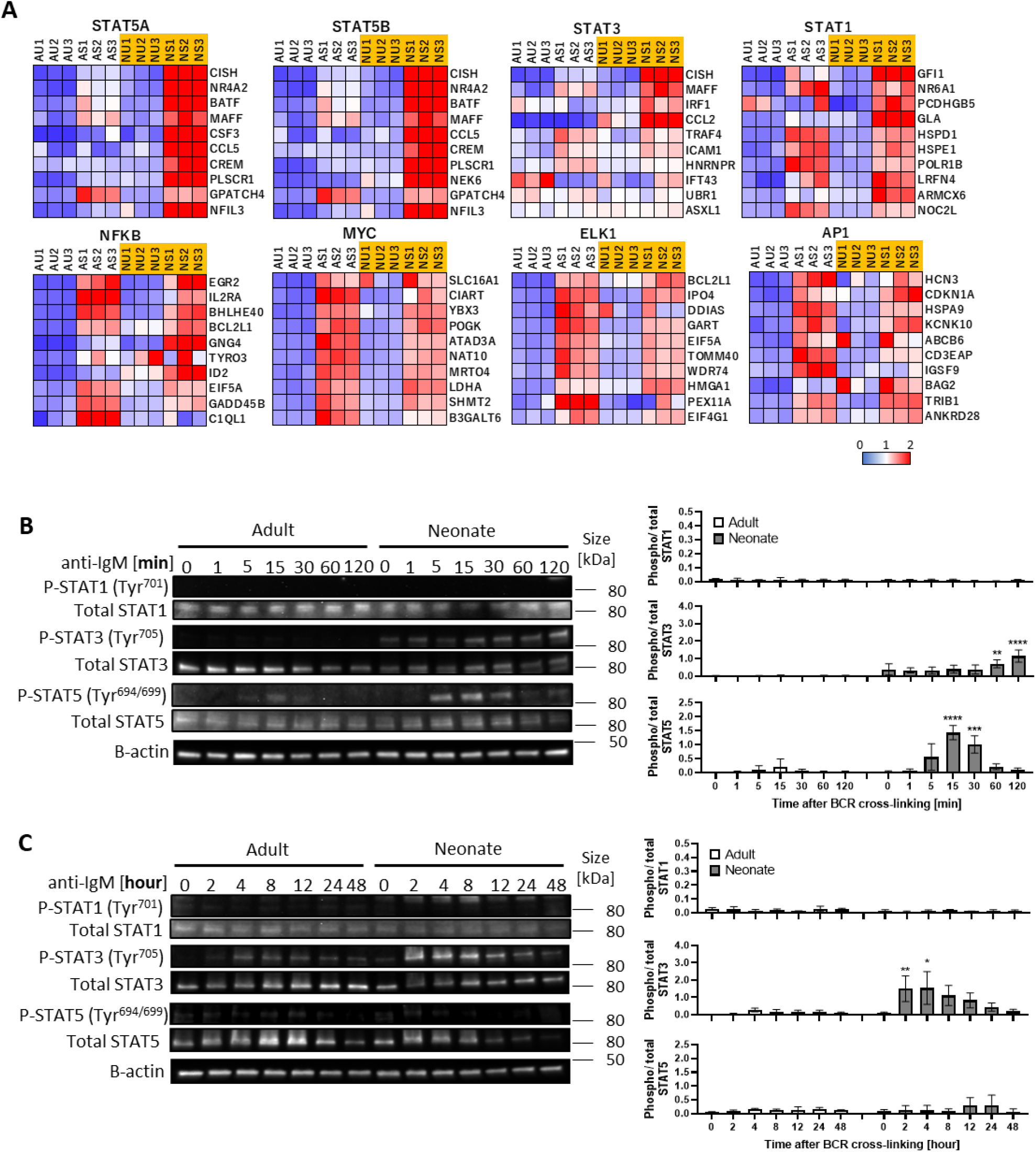
STAT3 and STAT5 are uniquely activated in neonatal B cells following BCR cross-linking. In all experiments splenic CD19^+^CD43^-^ B cells were isolated and stimulated with 10 μg/mL F(ab’)2 fragments of anti-IgM antibodies to engage BCR under different conditions. **(A)** RNA-seq analysis was performed on CD19^+^CD43^-^ cells stimulated with anti-IgM antibodies for 7 h. Total RNA was isolated for regular RNA sequencing. Gene set enrichment analysis (GSEA) was performed using the hallmark gene sets in C3 transcription factor targets (TFT). Heat maps show top 10 genes in selected TFs. AU: adult unstimulated; AS: adult stimulated; NU: neonate unstimulated; NS: neonate stimulated. (**B and C**) CD19^+^CD43^-^ cells were stimulated with anti-IgM antibodies for the indicated duration, and whole cell extracts were collected for immunoblot analysis of STAT1, STAT3 and STAT5. Data are shown as the mean ± s.d. of three independent experiments. *P* values were calculated using one-way ANOVA with a Dunnett’s multiple comparisons test (\**P*<0.05, \*\**P*<0.01, *** *P*<0.001, and **** *P*<0.0001).

### STAT3 and STAT5 are involved in neonatal BCR induced IL-10 production

We next asked whether the differential activation of STAT3 and STAT5 in BCR-stimulated adult and neonatal CD19^+^CD43^-^ cells help explain elevated IL-10 production from neonatal CD19^+^CD43^-^ cells. Although the involvement of STAT5 is not clear in IL-10 production, STAT3 has been shown to function as a promotor of IL-10 production in several types of cells, including human B cells (*40–43*). To assess the participation of these signaling molecules in IL-10 production, we used chemical inhibitors known to ablate STAT3 and STAT5 activation (*44, 45*). We found that both S3I-201 (STAT3 inhibitor) and Pimozide (STAT5 inhibitor) inhibited BCR-induced *Il10* mRNA expression and IL-10 protein production (Fig. 3, A and B). To gain further insight into the activation of STAT3 and STAT5 downstream of BCR, we focused on Janus-activated kinases (JAKs) because STAT proteins are activated by JAKs in cytokine receptor signaling pathways (*46*). We selected the pan-JAK inhibitor Pyridone 6 (*47*) to test the involvement of STAT3 and STAT5. We first verified the inhibitory activity of Pyridone 6 in B cells, which effectively inhibited STAT3 and STAT5 phosphorylations induced by IL-6 and IL-21, respectively (fig. S7A). When neonatal B cells were stimulated through BCR in the presence of Pyridone 6, STAT3 phosphorylation, but not STAT5 phosphorylation, was completely suppressed (Fig. 3C). Similarly, as previously observed (*48*), specific JAK2 inhibitor AG490 failed to inhibit STAT5 phosphorylation (fig. S7B). While neonatal BCR-induced STAT5 activation was JAK-independent, it was inhibited by Ibrutinib (Bruton’s tyrosine kinase (BTK) inhibitor) and Staurosporine (protein kinase C (PKC) inhibitor) as shown previously (*49, 50*) (fig. S7B). These results suggested that whereas BCR induced STAT3 phosphorylation relied on JAKs, STAT5 is activated in a JAK-independent and BTK/PKC dependent fashion.

**Figure 3.**
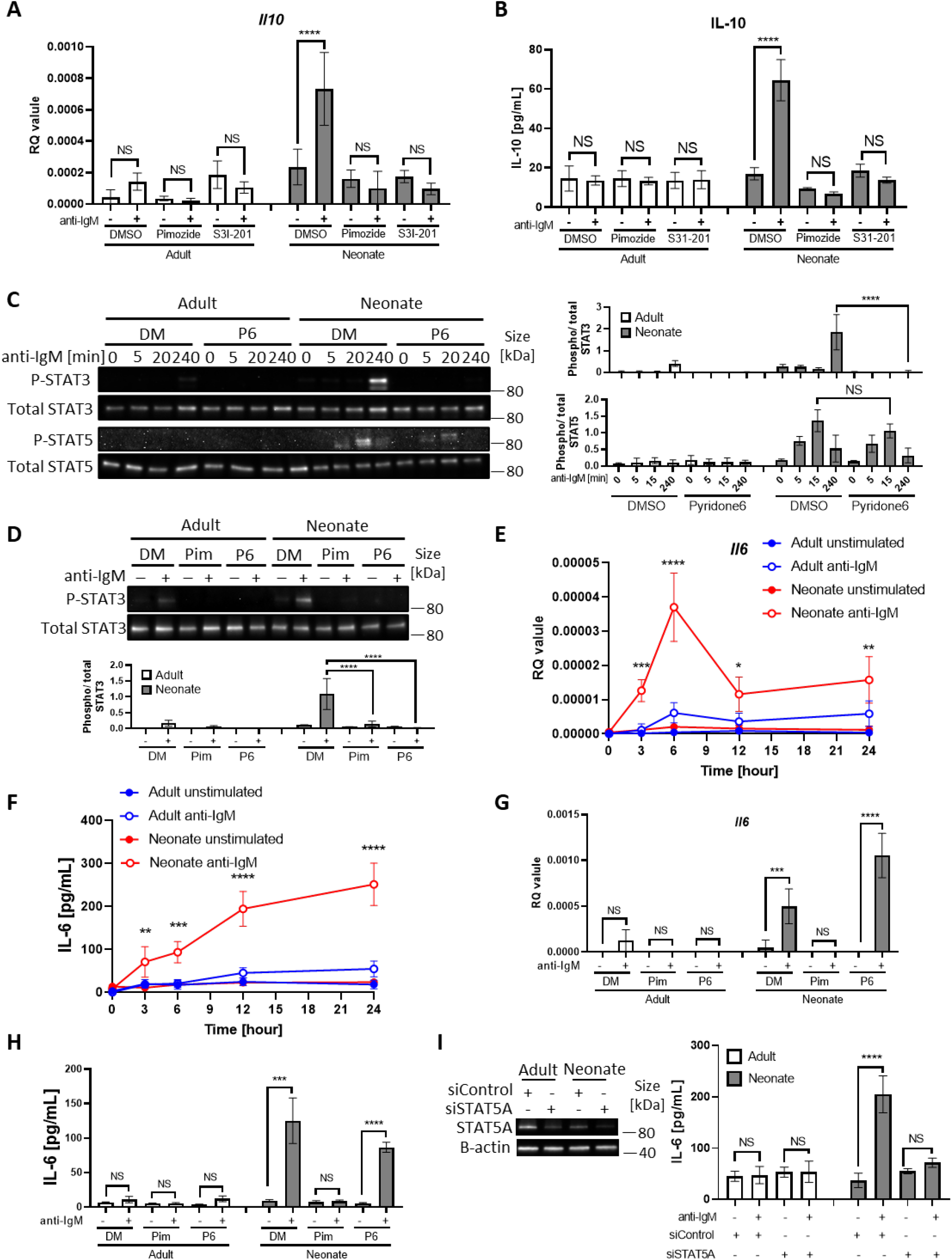
STAT3 is activated by autocrine IL-6 in a STAT5-dependent manner. In all experiments splenic CD19^+^CD43^-^ B cells were isolated and stimulated with 10 μg/mL F(ab’)2 fragments of anti-IgM antibodies to engage BCR under different conditions. **(A and B)** CD19^+^CD43^-^ cells were pre-treated with DMSO, 20 μM Pimozide, or 100 μM S3I-201 for 1 h and then incubated in the absence or presence of anti-IgM antibodies for 7 h (*n* = 3). *Il10* mRNA expression was determined by RT-qPCR (A) (*n* = 3) and the amount of secreted IL-10 was determined by ELISA (B) (*n* = 3). **(C)** CD19^+^CD43^-^ cells were pre-treated with DMSO (DM) or 1 μM Pyridone 6 (P6) for 1 h prior to stimulation with anti-IgM antibodies for the indicated duration and changes in STAT3 (Tyr^705^) and STAT5 (Tyr^694/699^) phosphorylations were detected in Western blot analysis. **(D)** CD19^+^CD43^-^ cells were pre-treated with DMSO, 20 μM Pimozide (Pim), or 1 μM Pyridone 6 (P6) for 1 h and then incubated in the absence or presence of anti-IgM antibodies for 4 h and changes in STAT3 (Tyr^705^) phosphorylation were detected in Western blot analysis. **(E and F)** CD19^+^CD43^-^ cells were incubated with anti-IgM antibodies for the indicated duration and *Il6* mRNA expression was determined by RT-qPCR (E) (*n* = 3) and the amount of secreted IL-6 was measured by ELISA (F) (*n* = 3). *P* values (Adult anti-IgM vs Neonate anti-IgM) were calculated using two-way ANOVA. **(G and H)** CD19^+^CD43^-^ cells were pre-treated with DMSO, 20 μM Pimozide, or 1 μM Pyridone 6 for 1 h and then incubated in the absence or presence of anti-IgM antibodies for 4 h and *Il6* mRNA expression was determined by RT-qPCR (G) (*n* = 3) and the amount of secreted IL-6 was determined by ELISA (H) (*n* = 3). **(I)** CD19^+^CD43^-^ cells were transfected with siRNA targeting *STAT5A* (siSTAT5A) or non-targeting control siRNA (siControl) for 48 h, and then incubated in the absence or presence of anti-IgM antibodies for 4 h. The amount of STAT5A was examined by immunoblot analysis. The amount of secreted IL-6 was determined by ELISA (Right) (*n* = 3). Unless otherwise is indicated, *P* values were calculated using one-way ANOVA with a Dunnett’s multiple comparisons test (\**P*<0.05, \*\**P*<0.01, \*\*\**P*<0.001, and\*\*\*\**P*<0.0001). No significant difference; NS. In all experiments data are shown as the mean ± s.d. of three independent experiments.

### Neonatal BCRs indirectly activate STAT3 via STAT5-dependent autocrine IL-6

Differences in the phosphorylation kinetics (Fig. 2, B and C) and the dependency on JAKs (Fig. 3C) between STAT3 and STAT5 in CD19^+^CD43^-^ cells led us to hypothesize that STAT5 had a role in mediating neonatal BCR signaling to STAT3 in a JAK-dependent manner. To test this hypothesis, we investigated whether the STAT5 inhibitor Pimozide would block STAT3 phosphorylation. Supporting this hypothesis and suggesting that STAT5 functions as an upstream mediator of STAT3, Pimozide suppressed neonatal BCR-induced STAT3 phosphorylation (Fig. 3D). Next, we sought to elucidate how STAT5 activates STAT3 in a JAK-dependent fashion. As JAKs are activated by cytokine receptors and several studies proposed autocrine IL-6 signaling in tumor cell lines (*51–55*), we envisaged an autocrine activity of IL-6 produced by BCR-stimulated neonatal CD19^+^CD43^-^ B cells. Indeed, we found significantly higher *Il6* mRNA expression and IL-6 protein production following BCR cross-linking in neonatal CD19^+^CD43^-^ B cells compared to those in adult counterparts (Fig. 3, E and F). Higher IL-6 production in neonatal B cells following BCR cross-linking was also observed by flow cytometry (fig. S7C). While BCR-mediated signaling alone is not sufficient to induce IL-6 in adult B cells (*56, 57*), studies in macrophages and cell lines have shown that STAT5 can contribute to IL-6 promoter activity (*51, 58*). To test whether the IL-6 production by neonatal B cells was also governed by STAT5 activation, we stimulated cells in the presence of the STAT5 inhibitor, Pimozide. Indeed, Pimozide abrogated both *Il6* mRNA expression and IL-6 protein production following BCR engagement, whereas the pan-JAK inhibitor Pyridone 6 did not (Fig. 3, G and H). This result was reproduced by using small interfering RNA targeting STAT5, which also suppressed BCR-induced IL-6 production (Fig. 3I). These findings indicated that STAT5 has an essential role in the production of IL-6 by BCR-activated neonatal CD19^+^CD43^-^ cells.

Next, we sought to investigate whether the STAT3 phosphorylation in BCR-activated neonatal cells was due to the IL-6 secreted from CD19^+^CD43^-^ cells. IL-6 signaling is mediated by membrane-bound or soluble IL-6 receptor alpha (IL-6Rα) and a transmembrane protein gp130 (*59*). Only after forming a complex with IL-6, the IL-6Rα associates with gp130 and transduces signals for JAK-STAT3 pathway (*60, 61*). Suggesting a role for IL-6 in STAT3 activation, blockade of IL-6 signaling with anti-IL-6Rα monoclonal antibody (15A7) reduced neonatal BCR-induced STAT3 phosphorylation, but not STAT5 phosphorylation (Fig. 4A). Similarly, the gp130 inhibitor SC144 abrogated STAT3 phosphorylation following BCR engagement (fig. S8). While the blocking experiments indicated that IL-6 produced in response to BCR-engagement was responsible for STAT3 activation, STAT3 is also activated by IL-10 signaling (*62*) (fig. S9A). Since BCR stimulation also induces IL-10 production in neonatal B cells, we tested the involvement of IL-10 in STAT3 phosphorylation by using anti-IL-10R blocking monoclonal antibody. We found that anti-IL-10R antibody did not reduce neonatal BCR-induced STAT3 phosphorylation while anti-IL-6Rα antibody did (fig. S9B). These results suggest that neonatal CD19^+^CD43^-^ cells produced IL-6 following BCR cross-linking in a STAT5-dependent manner, and subsequently the secreted IL-6 activated STAT3 via the IL-6R-gp130 pathway in an autocrine/paracrine fashion.

**Figure 4.**
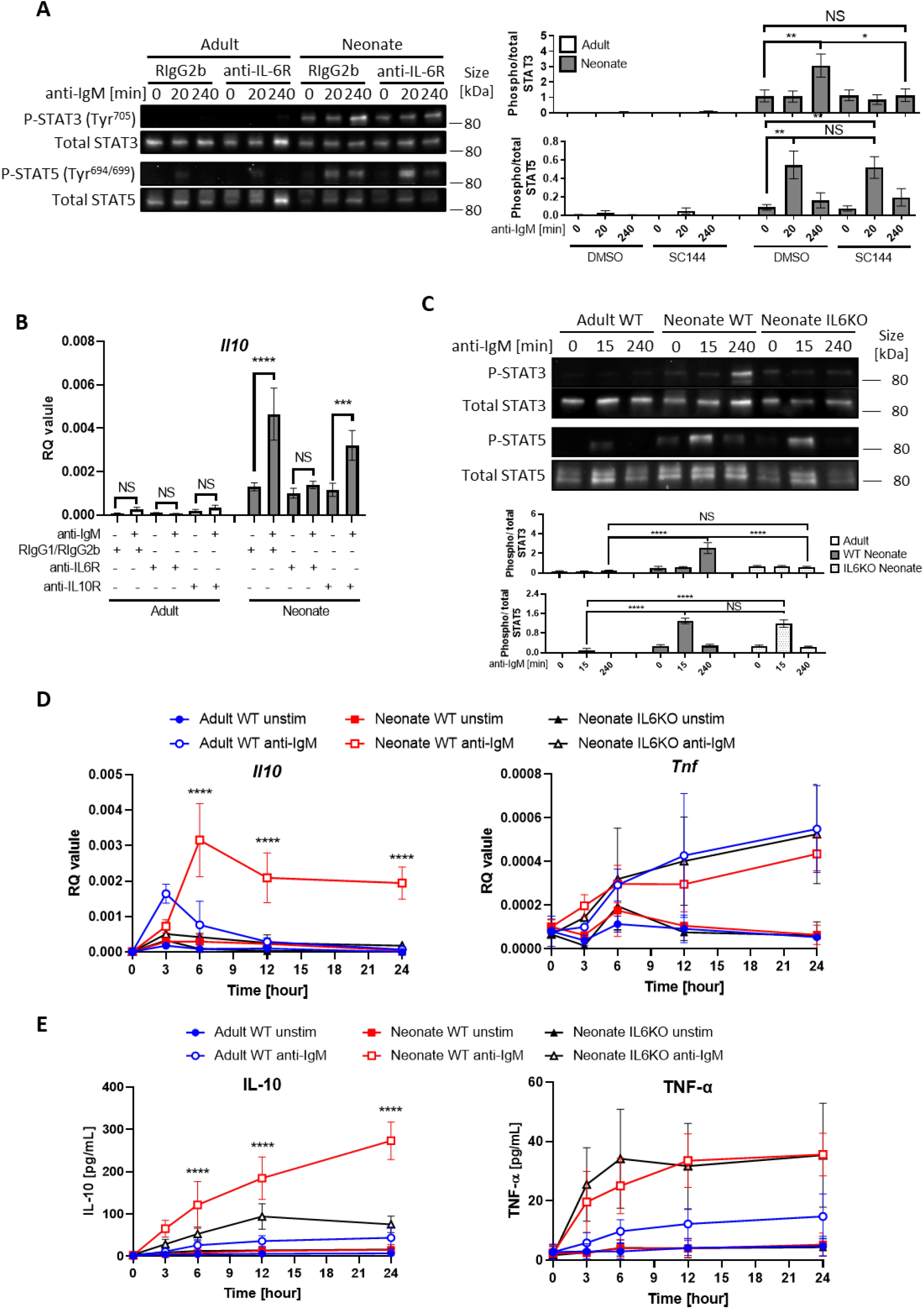
Autocrine IL-6 activates IL-10 expression in a STAT3-dependent manner. In all experiments splenic CD19^+^CD43^-^ B cells were isolated and stimulated with 10 μg/mL F(ab’)2 fragments of anti-IgM antibodies to engage BCR under different conditions. **(A)** CD19^+^CD43^-^ cells were stimulated with anti-IgM antibodies for the indicated duration and changes in STAT3 (Tyr^705^) and STAT5 (Tyr^694/699^) phosphorylations were detected in Western blot analysis. **(B)** CD19^+^CD43^-^ cells were pre-treated with 10 μg/mL isotype control antibodies (Rat IgG1/Rat IgG2b), anti-IL-6R antibody or anti-IL-10R antibody for 1 h prior to stimulation with of anti-IgM antibodies for 7 h and *Il10* mRNA expression was determined by RT-qPCR (*n* = 3). **(C)** CD19^+^CD43^-^ cells, isolated from wild-type (WT) adult, WT neonate, and IL-6-deficient (IL6KO) neonate, were stimulated with anti-IgM antibodies for the indicated duration and changes in STAT3 (Tyr^705^) and STAT5 (Tyr^694/699^) phosphorylations were detected in Western blot analysis. **(D and E)** CD19^+^CD43^-^ cells were incubated with anti-IgM antibodies for the indicated duration. *Il10* and *Tnf* mRNA expression were determined by RT-qPCR (D) (*n* = 3) and the amounts of secreted IL-10 and TNF-α were determined by ELISA (E) (*n* = 3). *P* values (Neonate WT anti-IgM vs Neonate IL6KO anti-IgM) were calculated using two-way ANOVA (\*\*\*\**P*<0.0001). Unless otherwise indicated, in other experiments *P* values were calculated using one-way ANOVA with a Dunnett’s multiple comparisons test (\**P*<0.05, \*\**P*<0.01, \*\*\**P*<0.001, and\*\*\*\**P*<0.0001). No significant difference; NS. In all experiments data are shown as the mean ± s.d. of three independent experiments.

### Neonatal B cells produce IL-10 in an autocrine fashion by IL-6-activated STAT3

The fact that both STAT3 and STAT5 inhibitors blocked *Il10* mRNA expression and IL-10 protein production from neonatal CD19^+^CD43^-^ cells (Fig. 3, A and B) and the IL-6 produced by BCR-stimulated CD19^+^CD43^-^ cells was responsible for the activation of STAT3 (Fig. 4A) suggested that IL-10 might be produced in response to IL-6. To begin addressing this question, we first investigated whether the production of IL-10 was dependent on IL-6 signaling in CD19^+^CD43^-^ cells. We found that blocking with antibodies against IL-6Rα, but not with anti-IL-10R, totally abrogated *Il10* mRNA expression seven hours after BCR stimulation (Fig. 4B). To further elucidate this question, we evaluated responses in splenic CD19^+^CD43^-^ B cells from IL-6-deficient neonatal mice. We began by assessing the activation of STATs in IL-6 knock out (KO) neonatal splenic CD19^+^CD43^-^ cells. Further underscoring the dependency of BCR-induced STAT3 activation on IL-6 production, BCR cross-linking did not induce STAT3 phosphorylation in CD19^+^CD43^-^ cells from IL-6 KO neonatal mice, whereas STAT5 phosphorylation was comparable between wild-type and IL-6 KO strains (Fig. 4C). We excluded the possibility of a defect in STAT3 activation in IL-6-deficient CD19^+^CD43^-^ cells because recombinant IL-6 induced the phosphorylation of STAT3 in IL-6 KO neonatal CD19^+^CD43^-^ cells (fig. S9C).

Next, we assessed the cytokine levels in IL-6-deficient neonatal B cells following BCR engagement. Both *Il10* gene expression and IL-10 protein levels were significantly reduced in IL-6 KO neonatal CD19^+^CD43^-^ cells compared to wild-type counterparts (Fig. 4, D and E). IL-6 deficiency did not create an overall suppressive state because both *Tnf* gene expression and TNF-α protein levels were comparable between wild-type and IL-6-deficient neonatal CD19^+^CD43^-^ cells (Fig. 4, D and E). Notably, despite the significant reduction in IL-10 levels, there were residual gene and protein expression in CD19^+^CD43^-^ cells from IL-6 KO mice. To test whether other cytokines also contribute to IL-10 production, we focused on IL-1β because IL-1β is shown to enhance IL-10 production by CD40-activated B cells (*15*). While neonatal CD19^+^CD43^-^ cells significantly increased *Il1b* gene expression following BCR cross-linking (fig. S10A), blocking IL-1β signaling using a neutralizing antibody failed to reduce IL-10 production by IL-6-deficient neonatal CD19^+^CD43^-^ B cells (fig. S10, B and C). In addition, biologically active recombinant IL-1β (fig. S10D) did not induce IL-10 while recombinant IL-6 did (fig. S10E). The IL-12 family member cytokine IL-35 has been shown to induce Breg differentiation and IL-10 production (*63*). The autocrine effect of IL-35 on IL-10 production by Bregs has been considered since this cytokine is secreted by Bregs as well as Tregs (*64*). To assess whether IL-35 may be involved in IL-10 secretion, we measured IL-35 production from anti-IgM stimulated neonatal B cells. Ruling out its involvement, IL-35 production was not observed following BCR cross-linking (fig. S11). These results suggested that molecule(s) other than IL-1β and IL-35 are likely responsible for the production of residual IL-10 from BCR-stimulated IL-6 KO neonatal CD19^+^CD43^-^ cells.

In addition to the elevated IL-6 production, flow cytometry assessment revealed that neonatal B cells expressed higher levels of surface IL-6Rα compared to their adult counterpart (Fig. 5A). The expression level of gp130 was comparable between the two age groups (fig. S12). To test whether higher IL-6Rα expression translated into enhanced IL-6 activity, we incubated purified CD19^+^CD43^-^ population in the presence of increasing concentrations of recombinant IL-6 and measured P-STAT3 in Western blot analysis. We detected P-STAT3 in neonatal cells with as little as 0.04 ng/mL of IL-6 whereas approximately 62 times more IL-6 (2.5 ng/mL) was needed to induce the phosphorylation of STAT3 in adult cells (Fig. 5B). Quantification of the IL-6 activity by computing half maximal effective concentration (EC_50_) indicated that the EC_50_ for neonatal B cells (0.98 ± 0.41 ng/mL) was significantly lower than that for adult counterparts (4.37 ± 1.27 ng/mL). Moreover, the elevated neonatal B cell P-STAT3 activity translated into increased IL-10 production in IL-6-stimulated neonatal CD19^+^CD43^-^ cells (Fig. 5C).

**Figure 5.**
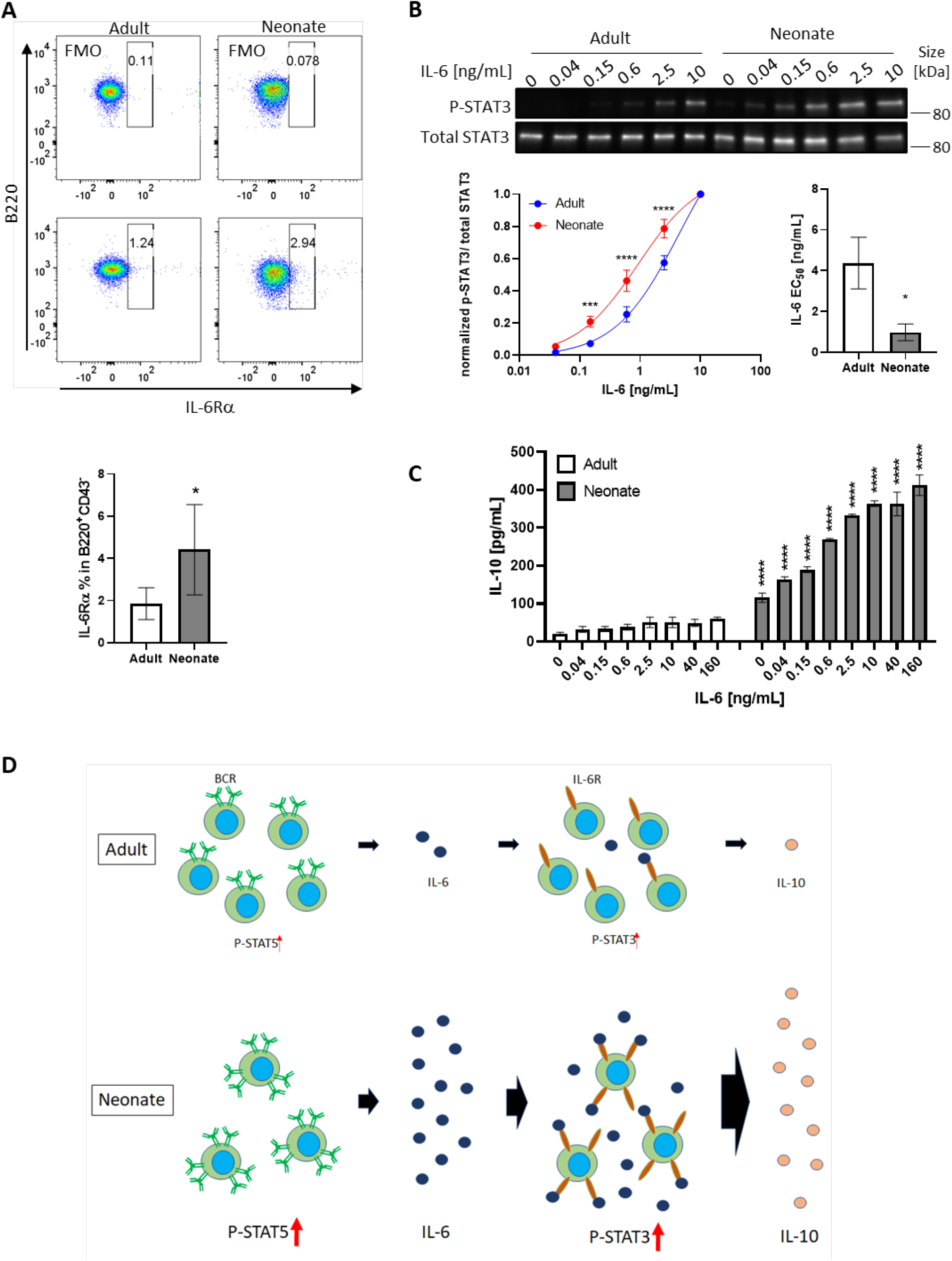
Neonatal BCRs activate STAT5 in PKC/FAK/Rac1/Syk and Btk signaling axis. In all experiments isolated splenic CD19^+^CD43^-^ B cells were used. **(A)** CD19^+^CD43^-^ cells were analyzed for surface levels of IL-6Rα (*n* = 3). Data are shown as the mean ± s.d. of three independent experiments. *P* values versus adult counterparts were calculated using two-tailed unpaired *t*-tests (\**P*<0.05). **(B)** CD19^+^CD43^-^ cells were stimulated with recombinant IL-6 at indicated concentrations for 30 min. Changes in STAT3 (Tyr^705^) phosphorylation were detected in Western blot analysis. Data are representative of three independent experiments. Data are shown as the mean ± s.d. of normalized ratio of P-STAT3 to total STAT3 in three independent experiments. *P* values versus adult counterparts were calculated using two-way ANOVA with a Sidak’s multiple comparisons test for the dose-response curve (\*\*\**P*<0.001 and \*\*\*\**P*<0.0001) and two-tailed Student’s *t*-test for the IL-6 EC50 comparison (\**P*<0.05). **(C)** CD19+CD43-cells were stimulated with increasing concentrations of recombinant IL-6 for 48 h. The amount of IL-10 secreted was determined by ELISA (n = 3). Data are shown as the mean ± s.d. of two independent experiments. *P* values versus adult counterparts were calculated using one-way ANOVA with a Dunnett’s multiple comparisons test (****P<0.0001). **(D)** Schematic representation of cellular events leading to enhanced IL-10 production from neonatal CD19^+^CD43^-^ B cells following BCR cross-linking is shown.

Finally, we wanted to examine whether IL-6 was also involved in the production of IL-10 in response to toll-like receptor (TLR) ligands because neonatal B cells have also been shown to produce elevated levels of IL-10 in response to TLR ligand stimulation (*65–67*). As reported previously, TLR9 and TLR4 ligands induced high levels of IL-10 from wild-type neonatal mouse CD19^+^CD43^-^ cells (fig. S13). IL-10 secreted by IL-6-deficient neonatal CD19^+^CD43^-^ cells were comparable to that from wild-type neonatal CD19^+^CD43^-^ cells in response to CpG and lipopolysaccharide (LPS), suggesting that TLR signaling-induced IL-10 production was not dependent on IL-6. Taken together, these results revealed a mechanism whereby neonatal CD19^+^CD43^-^ cell BCR-STAT5 axis-induces IL-6 which in turn activates STAT3 in autocrine/paracrine fashion, leading to IL-10 production (Fig. 5D).

### Neonatal BCR activates STAT5 via protein kinase C, focal adhesion kinase, and Rac1

We next sought to elucidate how neonatal BCRs activate STAT5 in a JAK-independent manner (Fig. 3C). We focused on the involvement of protein kinase C (PKC), focal adhesion kinase (FAK) and Rac1 because PKC was suggested to play a role in BCR-triggered STAT5 activation (*49*) and autophosphorylation of FAK and Rac1 mediates STAT5 activation in oncogenic cells (*68*). Moreover, PKC has been shown to promote FAK phosphorylation (*69–71*) and to activate Rac1 (*72, 73*). We found that BCR cross-linking increased the phosphorylation of PKC in neonatal CD19^+^CD43^-^ cells, but not in adult cells (Fig. 6A). We assessed the two phosphorylation sites of FAK: an autophosphorylation site (Tyr^397^) that functions as a Src homology 2 (SH2) binding site and the kinase domain (Tyr^567^) that promotes FAK catalytic activity (*74*). FAK was constitutively phosphorylated at Tyr^397^ in neonatal cells and this phosphorylation increased further after stimulation, whereas BCR cross-linking triggered a comparable increase in phosphorylation at Tyr^567^ in adults and neonates (Fig. 6A and fig. S14), suggesting that the FAK catalytic activity did not lead to the unique STAT5 phosphorylation in neonates. Confirming their upstream role in controlling STAT5 activation, both the PKC inhibitor staurosporine and the FAK inhibitor 14 (F-14) inhibited neonatal BCR-induced STAT5 phosphorylation (Fig. 6B). Moreover, *Il6* gene expression in response to neonatal BCR engagement was impeded by these inhibitors as well as the STAT5 inhibitor (Fig. 6C). Also, the Rac1 inhibitor, NSC23766 inhibited STAT5 phosphorylation (Fig. 6D) and *Il6* gene expression (Fig. 6E) following BCR cross-linking. Having established that PKC governs BCR-induced STAT5 phosphorylation, we tested whether the specific PKC activator phorbol 12-myristate 13-acetate (PMA) also triggered STAT5 phosphorylation. As expected, PKC was phosphorylated in both adult and neonatal CD19^+^CD43^-^ cells after PMA stimulation (fig. S15A). Also, PMA induced the phosphorylation of p65 and IκBα in both neonatal and adult CD19^+^CD43^-^ cells (fig. S15A). However, only neonatal B cells manifested STAT5 phosphorylation in response to PMA (fig. S15A). Finally, by using specific inhibitors, we found that spleen tyrosine kinase (Syk) and Btk control BCR-induced STAT5 phosphorylation (fig. S15B). Together, these results unveiled the upstream pathways involving Syk, Btk, PKC, FAK and Rac1 in BCR-induced *Il6* gene expression via STAT5 phosphorylation.

**Figure 6.**
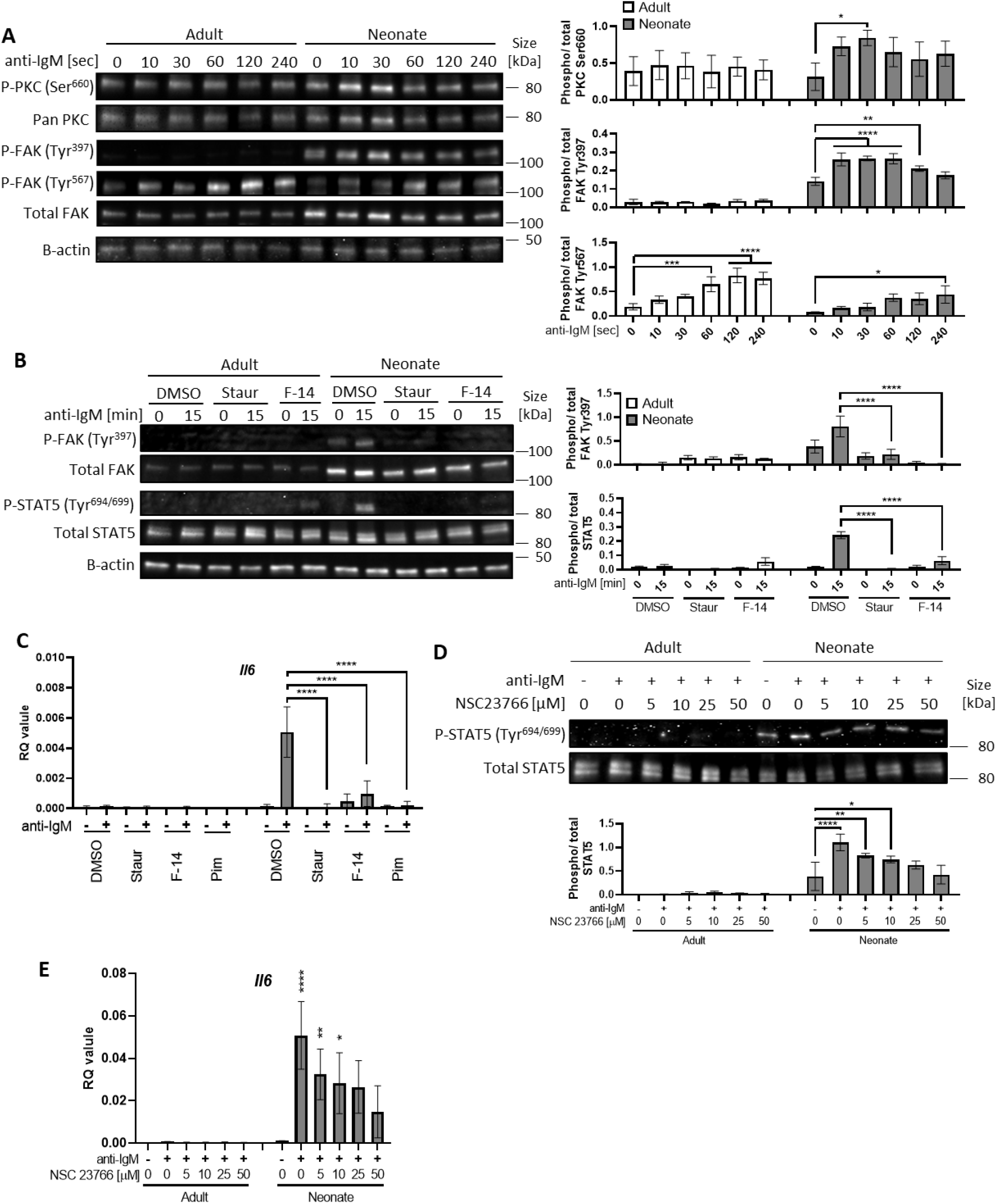
In all experiments splenic CD19^+^CD43^-^ B cells were isolated and stimulated with 10 μg/mL F(ab’)_2_ fragments of anti-IgM antibodies to engage BCR under different conditions. **(A)** CD19^+^CD43^-^ cells were stimulated with anti-IgM antibodies for the indicated duration and changes in PKC (Ser^660^), FAK (Tyr^397^) and FAK (Tyr^567^) phosphorylations were detected in Western blot analysis. **(B)** CD19^+^CD43^-^ cells were pre-treated with DMSO, 20 nM PKC inhibitor Staurosporine (Staur), or 10 μM FAK inhibitor 14 (F-14) for 1 h and then stimulated with anti-IgM antibodies for the indicated duration. Changes in FAK (Tyr^397^) and STAT5 (Tyr^694/699^) phosphorylations were detected in Western blot analysis. **(C)** Isolated CD19^+^CD43^-^ B cells were pre-treated with DMSO, 20 nM Staurosporine, or 10 μM F-14, or 20 μM Pimozide for 1 h prior to incubation in the absence or presence of anti-IgM antibodies for 4 h and *Il6* mRNA expression was determined by RT-qPCR (*n* = 3). **(D)** CD19^+^CD43^-^ cells were pre-treated with Rac1 inhibitor NSC23766 for 1 h and then stimulated with anti-IgM antibodies for 15 min. Changes in STAT5 (Tyr^694/699^) phosphorylation was detected in Western blot analysis. **(E)** Isolated CD19^+^CD43^-^ B cells were pre-treated with NSC23766 for 1 h prior to incubation in the absence or presence of anti-IgM antibodies for 20 h and *Il6* mRNA expression was determined by RT-qPCR (*n* = 3). In all experiment data shown as the mean ± s.d. of three independent experiments. *P* values were calculated using one-way ANOVA with a Dunnett’s multiple comparisons test (\**P*<0.05, \*\**P*<0.01, \*\*\**P*<0.001, and \*\*\*\**P*<0.0001).

### IL-6 is not responsible for IL-10 production by CD43-expressing B-1 subset in neonatal mouse

Using a range of experiments, we demonstrated that BCR-crosslinking of CD19^+^CD43^-^ B cells lead to IL-6 production and this IL-6 is responsible for IL-10 secretion from CD19^+^CD43^-^ B cells (Fig. 4B, D, and E). Among the neonatal splenic B cells, the highest IL-10-producing subset is the CD43-expressing B-1 subset (Fig. 1B). To determine whether IL-6 is also responsible for IL-10 production by this subset, we compared IL-10-producing B-1 subset from BCR-stimulated wild-type and IL-6 KO neonatal B cells. We found that the BCR cross-linking did not increase CD19^+^CD43^+^IL-10^+^ population in both wild-type and IL-6 KO cells (fig. S16). Thus, BCR-induced IL-6 does not promote the expression of IL-10 by B-1 population.

### IL-10 secreted from neonatal CD19^+^CD43^-^ cells suppress TNF-α production by macrophages

Bregs exert their immune suppressive functions via both IL-10-dependent and -independent mechanisms (*77*). IL-10 derived from Bregs has a major role in inhibiting inflammatory cytokine production by monocytes (*78, 79*). Neonatal B cells have been shown to restrict Th1 responses in vivo and ex vivo by suppressing myeloid cell functions in an IL-10-dependent manner (*25, 65, 80*). We next sought to determine whether neonatal BCR-induced autocrine IL-6 signaling has a role in the IL-10-dependent regulatory function of neonatal CD19^+^CD43^-^ cells. To this end, conditioned media (CM) were prepared from BCR-stimulated splenic CD19^+^CD43^-^ cells isolated from adult, wild-type neonatal (WT), and IL-6 KO neonatal mice and the inhibitory potential of the CM from each condition was tested by measuring TNF-α production from adult mouse peritoneal macrophages (fig. S17). We confirmed that CM from WT neonatal CD19^+^CD43^-^ cells contained the highest amount of IL-10 (Fig. 7A), whereas all three CMs contained low amount of TNF-α (Fig. 7B). When incubated with macrophages, unstimulated cell-CMs from all three groups failed to induce TNF-α production (Fig. 7C). In the presence of control antibody (Rat IgG1), CMs from anti-IgM-stimulated adult and IL-6 KO neonatal cells elicited significant and comparable increases in TNFα production (Fig. 7C). In sharp contrast, CM from anti-IgM-stimulated wild-type neonatal cells triggered a blunted TNF-α production from macrophages. To test whether the low TNF-α secretion with CM from wild-type neonate was due to elevated IL10 in the CM, we included blocking antibodies against IL-10R in the culture system. Suggesting a suppressive role for the elevated IL-10 levels in the CM from wild-type neonatal CD19^+^CD43^-^cells, IL-10R antibodies significantly increased the TNF-α production and restored its concentration to macrophages incubated with neonatal IL-6 KO CM. Since adult and IL-6 KO CM contained low IL-10 levels, addition of anti-IL-10R antibodies did not significantly alter the TNF-α production in macrophages incubated with these CMs. Thus, these results demonstrate a major role for autocrine IL-6 signaling in the suppressive effect of BCR-induced IL-10 from neonatal CD19^+^CD43^-^ B cells on macrophage inflammatory cytokine production.

**Figure 7.**
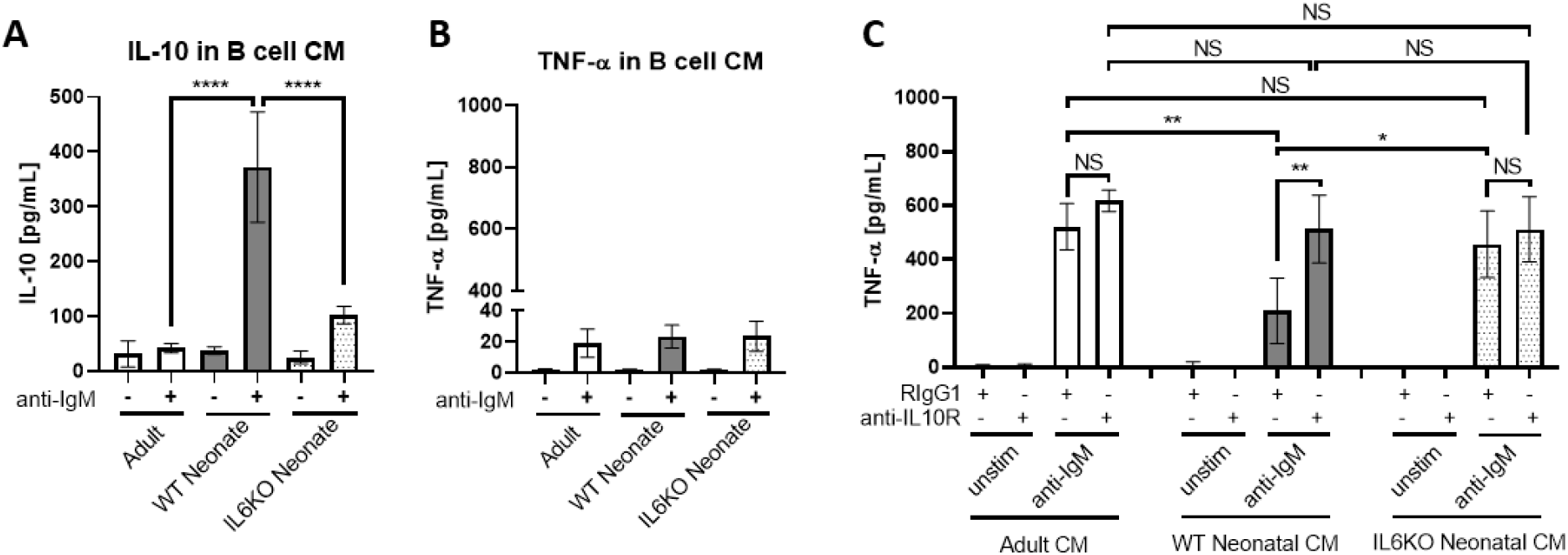
IL-10 secreted from neonatal CD19^+^CD43^-^ B cells suppress TNF-α secretion from macrophages. **(A and B)** Isolated splenic CD19^+^CD43^-^ B cells were stimulated with 10 μg/mL F(ab’)2 fragments of anti-IgM antibodies for 24 h. IL-10 (A) and TNF-α (B) levels in conditioned medium (CM) were determined by ELISA (*n* = 3). **(C)** Peritoneal macrophages isolated from adult mice were treated with 10 μg/mL anti-IL10R or isotype (Rat IgG1) control antibodies for 1h and then cultured in CM for 24 h. Levels of TNF-α secreted by macrophages cultured in CM were determined by ELISA (*n* = 3). Data are shown as the mean ± s.d. of three independent experiments. *P* values were calculated using one-way ANOVA with a Dunnett’s multiple comparisons test (\**P*<0.05, \*\**P*<0.01, and \*\*\*\**P*<0.0001).

## DISCUSSION

The propensity of neonatal splenic B cells to produce IL-10 with suppressive properties is well established (*13*). The increased IL-10 production by neonatal splenic B cells is primarily attributed to the B-1 cells, which constitute approximately 30% of B cells in neonatal spleen compared to less than 5% in adult spleen (*18*). Here, we described an additional subset of splenic B cells in neonates with higher IL-10 production potential than their adult counterparts. We found that BCR stimulation triggered IL-10 production by neonatal CD19^+^CD43^-^ B cells. In CD19^+^CD43^-^ cells, BCR stimulation led to STAT5 activation under the control of Syk, Btk, PKC, FAK and Rac1. Interestingly, BCR-induced STAT5 activation did not directly lead to IL-10 expression. Instead, the increase in IL-10 production was mostly dependent on STAT5-mediated IL-6 secretion, which promoted IL-10 secretion in an autocrine and paracrine fashion.

The IL-10 dependent suppression of inflammation by B10 cells is well documented in adult mouse autoimmune, allergy and infection models (*66, 81–84*). Despite the well-established requirement for antigen-mediated BCR signaling in B10 cell development, BCR-stimulation is not sufficient to induce the production of IL-10 from adult B10 cells (*26*). In fact, not only anti-IgM stimulation does not induce the secretion of IL-10 from adult B10 cells, but it also inhibits IL-10 production from LPS- and CD40L-stimulated B10 cells (*26*). Our results confirm the blunted IL-10 secretion from anti-IgM-stimulated adult B10 cells. Although neonatal B10 cells were reported previously by Yanaba and colleagues, the production of IL-10 from these cells in response to anti-IgM stimulation has not been studied (*26*). Our data show that neonatal CD19^+^CD43^-^ B cells are highly sensitive to BCR stimulation, but the secretion of IL-10 depends on the production of IL-6 from these cells rather than directly linking BCR to the transcription of *Il10* (fig. S18). In contrast to CD19^+^CD43^-^ B cells, B-1 cells secrete IL-10 spontaneously and the secretion of IL-10 is further increased following BCR-stimulation (*12*). The BCR-mediated IL-10 production from B-1 cells depends on p38 MAPK activity (*12*). The absence of IL-10 production directly after BCR stimulation in neonatal CD19^+^CD43^-^ B cells in our experiments is not unlikely to be due to the absence of p38 MAPK activity in these cells, because adult CD19^+^CD43^-^ B cells also failed to secrete IL-10 despite the phosphorylation of Akt, p38, JNK and ERK in response to BCR engagement. We found that the rapid IL-6 secretion following BCR engagement requires STAT5 activation (fig. S18). We also demonstrated that neonatal SH2 binding site (Tyr^397^) on FAK was constitutively phosphorylated, and this phosphorylation was required for STAT5 phosphorylation because the FAK inhibitor, F-14 which inhibits phosphorylation at Tyr^397^ effectively blocked BCR-induced STAT5 phosphorylation. These results indicated that FAK scaffold functions were essential to mediate BCR signaling to STAT5 in neonatal B cells as has been shown for FLT3 receptor signaling in oncogenic cells (*68*). A member of Rho family of GTPases, Rac1 has been shown to have a role in STAT5 activation in several types of cells (*85, 86*) and Rac1 functions as a downstream effector of FAK (*68*). Using specific inhibitors, we determined that Rac1, Syk, Btk, and PKC controlled neonatal BCR-induced STAT5 activation. Taken together, our data unveiled the signaling pathway mediated by Syk, Btk, PKC, FAK and Rac1 in the activation of STAT5 and the production of IL-6 in BCR stimulated neonatal CD19^+^CD43^-^ B cells (fig. S18).

Induction of IL-10 secretion from Breg cells by IL-6 has been shown previously by Rosser and colleagues in adult mice (*15*). However, in this study the authors have discovered that IL-6, as well as IL-1β, produced by macrophages in response to microbiota mediated stimuli act on splenic and lymph node Breg cells to produce IL-10. An important distinction between our findings in neonatal CD19^+^CD43^-^ B cells and those of Rosser and colleagues in adult Breg cells is that whereas microbiota-derived IL-6 and IL-1β required simultaneous CD40 signaling to induce IL-10 production from adult Breg cells, IL-6 alone was able to stimulate the production of IL-10 from neonatal CD19^+^CD43^-^ B cells. The independence of neonatal CD19^+^CD43^-^ B cells from CD40 signaling in IL-6-induced IL-10 secretion has in vivo relevance because neonatal B cells do not get activated through CD40 due to weak CD40L expression in neonatal T cells (*87, 88*). As was observed by Rosser and colleagues (*15*), we also found that recombinant IL-6 was not able to induce IL-10 production from adult CD19^+^CD43^-^ B cells. We think that the heightened sensitivity of neonatal B cells to IL-6 is likely due to the elevated expression of IL-6R on neonatal CD19^+^CD43^-^ B cells, which clearly rendered neonatal CD19^+^CD43^-^ B cells more sensitive to IL-6-mediated signaling. Higher IL-6 production by neonatal B cells following BCR engagement may have implications other than suppression through increased IL-10 secretion. The beneficial effect of IL-6 in the generation of adult T follicular helper (Tfh) cells, which help activate germinal center B cells to differentiate into plasma cells and memory B cells in response to immunization and infection is well recognized (*89, 90*). We have previously shown that, in contrast to adults, IL-6 suppresses vaccine response in neonatal mice by inhibiting Tfh generation and expanding the suppressive T follicular regulatory helper (Tfr) cells (*91*). Thus, in addition to stimulating IL-10 production, increased IL-6 secretion from BCR-stimulated neonatal B cells may be blunting Tfh generation.

The IL-10 produced by neonatal BCR-stimulated CD19^+^CD43^-^ B cells was functional because it inhibited TNF-α production by macrophages. We demonstrated that CM from anti-IgM-stimulated neonatal and adult B cells produced molecules that induced TNF-α production by macrophages, but this induction was blunted with the IL-10 in neonatal CM because the inhibitory anti-IL-10R antibody effectively restored the TNF-α secretion from macrophages. Moreover, there was no inhibition of TNF-α production from macrophages when CM from IL-6 KO B cells were used. In summary, our study revealed a detailed molecular and cellular mechanism involved in BCR-mediated IL-10 production by neonatal CD19^+^CD43^-^ B cells (fig. S18). This unique inhibitory pathway involving intermediate IL-6 secretion expands our understanding of the overall weak responses of neonates to vaccines and their susceptibility to infections.

## MATERIALS AND METHODS

### Mice

C57BL/6J mice and IL-6^-/-^ (B6.129S2-*Il6^tm1Kopf^*/J) mice were purchased from The Jackson Laboratory and maintained in local facilities. Two-to ten-month-old male and female were used in mating. All mice were fed regular chow in a pathogen-free environment. The mouse described in this study were performed in accordance with the US Food and Drug Administration/Center for Biologics Evaluation and Research Institutional Animal Care and Use Committee (permit 2002-31 and 2017-45).

### Cell isolation and culture

B lymphocytes were isolated from spleen tissues obtained from neonates (6-8 days old) and adult (8-10 weeks old) female mice. Total splenic B cells (CD19^+^) were purified using CD19 MicroBeads (130-121-301, Miltenyi Biotec, San Jose, CA), and CD19^+^CD43^-^ non-B-1 cells were purified using Mouse B Cell Isolation Kit (130-090-862, Miltenyi Biotec) which contains anti-CD43 antibody. The purity of isolated cells was assessed as B220^+^ and CD19^+^ double positive population with flow cytometry. Peritoneal macrophages were isolated from adult (8-10 weeks old) female mice using Peritoneum Macrophage Isolation Kit (130-110-434, Miltenyi Biotec). Isolated cells were cultured in complete RPMI+GlutaMAX (Thermo Fisher Scientific, Waltham, MA) supplemented with 10% fetal bovine serum (Life Technologies, Frederick, MD), 2 mM glutamine (Life Technologies), 100 U/mL penicillin and 100 μg/ml streptomycin (Life Technologies), 1mM sodium pyruvate (Life Technologies), 10 μM 2-mercaptoethanol (Sigma Aldrich, St Louis, MO), 20 mM HEPES (Life Technologies), and 1 mM MEM nonessential amino acids (Life Technologies). Cells were maintained in an incubator at 37°C, 5% CO2. Prior to stimulation, cells were treated with Pyridone 6 (Tocris Bioscience, Minneapolis, MN), Pimozide (Tocris Bioscience), S3I-201 (Sigma Aldrich), Staurosporine (Sigma Aldrich), FAK Inhibitor 14 (F-14) (R&D Systems, Minneapolis, MN), NSC23766 (R&D Systems), AG490 (Tocris Bioscience), SC144 (Tocris Bioscience), Syk Inhibitor (CAS622387-85-3, Sigma Aldrich), Ibrutinib (PCI-32765, Selleck Chemicals, Houston, TX), InVivoPlus rat IgG1 (HRPN, BioXcell, Lebanon, NH), InVivoMAb rat IgG2b (LTF-2, BioXcell), InVivo polyclonal Armenian hamster IgG (BioXcell), InVivoMAb anti-mouse IL-6R (15A7, BioXcell), InVivoPlus anti-mouse IL-10R (1B1.3A, BioXcell) or InVivoMAb anti-mouse/rat IL-1β (B122, BioXcell) for the indicated duration. Cells were stimulated with f(ab’)_2_ fragments of goat anti-mouse IgM antibody (eBioscience, San Diego, CA), recombinant mouse IL-6 (R&D Systems), recombinant mouse IL-21 (R&D Systems), recombinant mouse IL-10 (R&D Systems), recombinant mouse IL-1β (R&D Systems), CpG oligodeoxynucleotide 1826, or *Escherichia coli* lipopolysaccharide (Sigma Aldrich) for the indicated duration.

### Small interfering RNA (siRNA)

The following siRNAs were used in this study: Accell Mouse Stat5a (20850) siRNA-SMART pool (Dharmacon, Cambridge, UK) and Accell Non-targeting Pool (Dharmacon). Transfection was performed by following the manufacture’s instruction. Briefly, isolated cells were incubated with 1 μM Accell siRNA in Accell siRNA Delivery Media (Dharmacon) at 37°C, 5% CO_2_ for 48 h. Knockdown was assessed by Western blotting.

### RNA sequencing (RNA-seq)

Total RNA was extracted from isolated cells using the RNeasy Plus Mini kit (Qiagen, Germantown, MD) according to the manufacturer’s instruction. Briefly, isolated cells were lysed in RLT Plus buffer. Prior to RNA isolation, genomic DNA (gDNA) was removed using a gDNA eliminator. RNA was extracted after washing with RW1 buffer and RPE buffer. RNA samples were processed following the protocol for the Clontech (Mountain View, CA) SMART-Seq V4 Ultra Low Input RNA Preparation Kit (first-strand cDNA synthesis; full-length double strand cDNA amplification by LD-PCR; amplified cDNA purification and validation). The cDNAs were then further prepared using Illumina (San Diego, CA) Nextera DNA Library Preparation Kit (tagmentation; PCR amplification; clean-up and validation). Paired-end sequencing (100×2 cycles paired end reads) of multiplexed mRNA samples was carried out on an Illumina HiSeq 2500 sequencer. Fastq files obtained from the sequencer were generated using FDA HIVE platform v2.4 (*92*). Quality of sequence reads was inspected using HIVE’s native multi-QC tool. Sequence reads were mapped to GRCm38.p6 transcriptome (G CF_000001635.26) using HIVE’s native NGS aligner Hexagon (v2.4) (*93*). Gene-level feature counts were quantified using HIVE’s native feature mapping tool, HIVE Alignment Comparator (v2.4) and normalized to RPKM (*94*).

### Western Blot Analysis

Cultured cells were rinsed with ice-cold PBS with protease/phosphatase inhibitors (Thermo Fisher Scientific). Total cell lysates were prepared using RIPA buffer (Thermo Fisher Scientific) and protease/phosphatase inhibitors. The cell lysates were added into 4x Laemmli loading buffer (Bio Rad, Hercules, CA) and denatured at 95°C for 10 min. Protein samples were separated by tris-glycine denaturing SDS-PAGE (Bio Rad) and transferred onto nitrocellulose membranes (iBlot2, Invitrogen, Waltham, MA). Membranes were blocked with 5% bovine serum albumin or nonfat dry milk for 30 minutes, followed by incubation with primary antibodies overnight at 4°C and HRP-conjugated secondary antibodies for 1 hour. The following antibodies used in blotting were from Cell Signaling Technology (Danvers, MA): anti-phospho STAT1 Tyr^701^ (58D6), anti-STAT1 (42H3), anti-phospho STAT3 Tyr^706^ (D3A7), anti-STAT3 (79D7), anti-phospho STAT5 Tyr^694^ (D47E7), anti-STAT5 (D2O6Y), anti-beta-actin (D6A8), anti-phospho PKC (pan) betaII Ser^660^, anti-phospho FAK Tyr^397^, anti-FAK, anti-phospho Akt Ser^473^ (D9E), anti-Akt (pan) (C67E7), anti-phospho p38 MAPK Thr^180^/Tyr^181^ (D3F9), anti-p38 MAPK (D13E1), anti-phospho SAPK/JNK Thr^183^/Tyr^185^ (81E11), anti-SAPK/JNK, anti-phospho p44/42 MAPK (Erk1/2) Thr^202^/Tyr^204^ (D13.14.4E), anti-p44/42 MAPK (Erk1/2) (137F5), anti-phospho NF-κB p65 Ser^536^ (93H1), anti-NF-κB p65 (D14E12), anti-phospho IκBα Ser^32^ (14D4), anti-IκBα (44D4), anti-rabbit IgG, HRP-linked. Also, anti-PKC (A-3, Santa Cruz, Dallas TX), and anti-phospho FAK Tyr^576^ (2H74L24, Invitrogen) were used in Western blot analysis. Acquired images were analyzed using ImageJ (NIH) software.

### Flow Cytometry

For surface staining, cells were incubated with fluorochrome-labeled antibodies against surface proteins at 4 °C for 10 min, followed by 4’,6-diamino-2-phenylindole (DAPI) staining during a 5-min centrifugation. For all intracellular staining, cells were incubated in PBS containing Zombie UV^™^ Fixable Viability Dye (BioLegend, San Diego, CA) at 4 °C for 20 min.

Subsequently, for intracellular cytokine staining, cells were fixed/permeabilized with the Foxp3 staining buffer kit (Invitrogen) and then stained with fluorochrome-labeled antibodies against cytokines for 30 min at room temperature in Permeabilization Buffer (Invitrogen). For intracellular phosphorylated protein staining, cells were fixed with BD Cytofix™ (BD Biosciences) at 37 °C for 10 min and then permeabilized with ice-cold BD Phosflow™ Perm Buffer III (BD Biosciences) at 4 °C for 1 h. Cells were stained with fluorochrome-labeled antibodies against phosphorylated proteins for 30 min at room temperature in PBS containing 0.05% fetal calf serum. The following fluorochrome-labeled antibodies were from BioLegend: phycoerythrin (PE)-anti-mouse CD43 (S11), Pacific Blue-anti-mouse CD19 (6D5), fluorescein isothiocyanate (FITC)-anti-mouse CD19 (1D3/CD19), allophycocyanin (APC)-Cy7-anti-mouse CD19 (6D5), Brilliant Violet (BV) 510-anti-mouse/human CD45R/B220 (RA3-6B2), PE-anti-mouse CD126 (IL-6Rα chain) (D7715A7), FITC-anti-mouse IL-10 (JES5-16E3), and PE-anti-mouse IL-6 (MP5-20F3). Also, FAK (Phospho-Tyr^397^) (CF405M) (biorbyt, Cambridge, UK), FAK (FITC) (biorbyt), anti-phospho STAT3 Tyr^706^ (D3A7, Cell Signaling Technology), anti-phospho STAT5 Tyr^694^ (D47E7, Cell Signaling Technology), APC-anti-mouse CD130 (gp130) (KGP130, Invitrogen), and Alexa Fluor®488-Donkey anti-Rabbit IgG (H+L) (Jackson ImmunoResearch Inc.) were used in flow cytometry. Live cells were determined by forward Scatter (FSC), side scatter (SSC), DAPI staining or Zombie UV^™^ Fixable Viability Kit (BioLegend). Flow cytometric analysis of isolated B lymphocytes were performed using an LSR Fortessa flow cytometer (BD Biosciences) in the CBER Flow Cytometry Core Facility on the FDA White Oak campus (Silver Spring, MD). Data were analyzed using FlowJo software.

### Quantitative RT-PCR

To quantify gene expression, total RNA was extracted from isolated cells using the RNeasy Mini kit (Qiagen). Complementary DNA was synthesized from the extracted RNA using TaqMan™ Reverse Transcription Reagents (Applied Biosystems™, Bedford, MA). Quantitative PCR was performed on CFX96 Touch Real-Time PCR Detection System (Bio Rad) using TaqMan™ Gene Expression Master Mix (Applied Biosystems™) and TaqMan probes (Thermo Fisher Scientific). The following primers were used: *Gapdh* (Mm99999915_g1), *Il10*(Mm01288386_m1), *Il6* (Mm00446190_m1), *Tnf* (Mm00443258_m1), and *Il1b* (Mm00434228_m1). Delta Ct values were calculated with *Gapdh* and raised to the power of −2.

### Enzyme-linked immunosorbent assay (ELISA)

The levels of cytokines in cell culture medium were determined with BD OptEIA™ Mouse IL-6 ELISA set (BD Biosciences, Franklin Lakes, NJ), ELISA MAX™ Standard Set Mouse IL-10 (BioLegend), Mouse TNF-α DuoSet ELISA (R&D Systems), and LEGEND MAX™ Mouse IL-35 Heterodimer ELISA Kit (BioLegend) following manufacturer’s instructions.

### Statistical analyses

All statistical analyses were performed using Prism8 (GraphPad, San Diego, CA). Data were generally presented as mean ± standard deviation (s.d.). Two-tailed Student’s *t*-test, one-way analysis of variance (ANOVA) with a Dunnett’s multiple comparisons test and two-way ANOVA tests were performed to determine significance. *P* < 0.05 was considered statistically significant.

## Acknowledgments

We acknowledge the valuable technical support by FDA/CBER Veterinary Services and the Flow Cytometry Core Facility.

## Author contributions

Conceptualization: J.S. and M.A. Methodology: J.S., J.Y., C.C., W.W.W. and M.A. Investigation: J.S., J.Y. and M.A. Visualization: J.S. and M.A. Funding acquisition: M.A. Project administration: M.A. Supervision: M.A. Writing – original draft: J.S. and M.A. Writing – review & editing: J.S., J.Y., C.C., W.W.W. and M.A.

## Data and materials availability

All data needed to evaluate the conclusions in the paper are present in the paper or the Supplementary Materials.

