## Supplemental Figures for "B cell receptor induced IL-10 production from neonatal CD19+CD43- cells depends on STAT5 mediated IL-6 secretion"

**Figure S1**

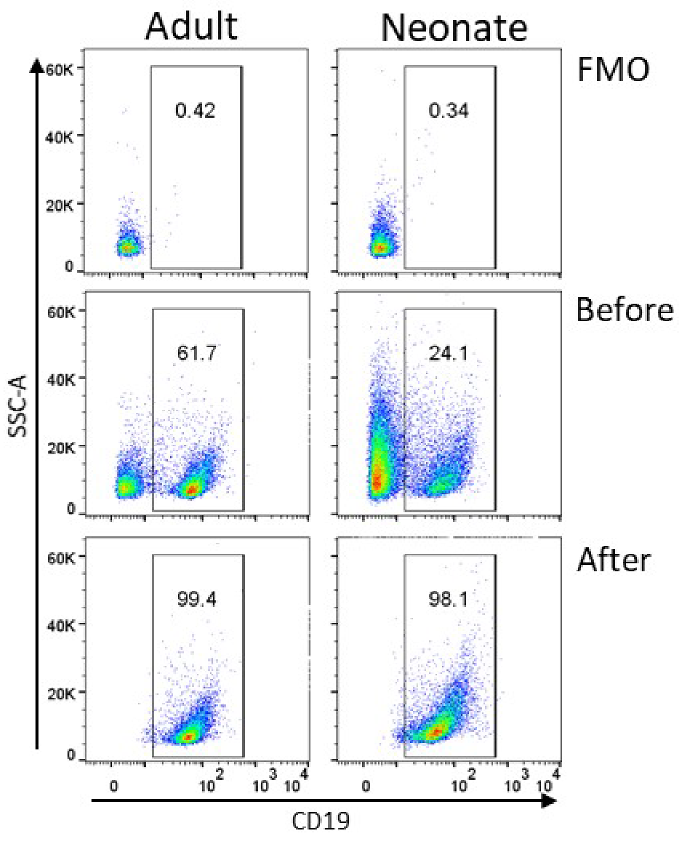


**Supplementary Figure S1. Isolation of splenic CD19$^+$ B cells.** Splenic CD19$^+$ B cells were isolated using CD19 MicroBeads.

**Figure S2**

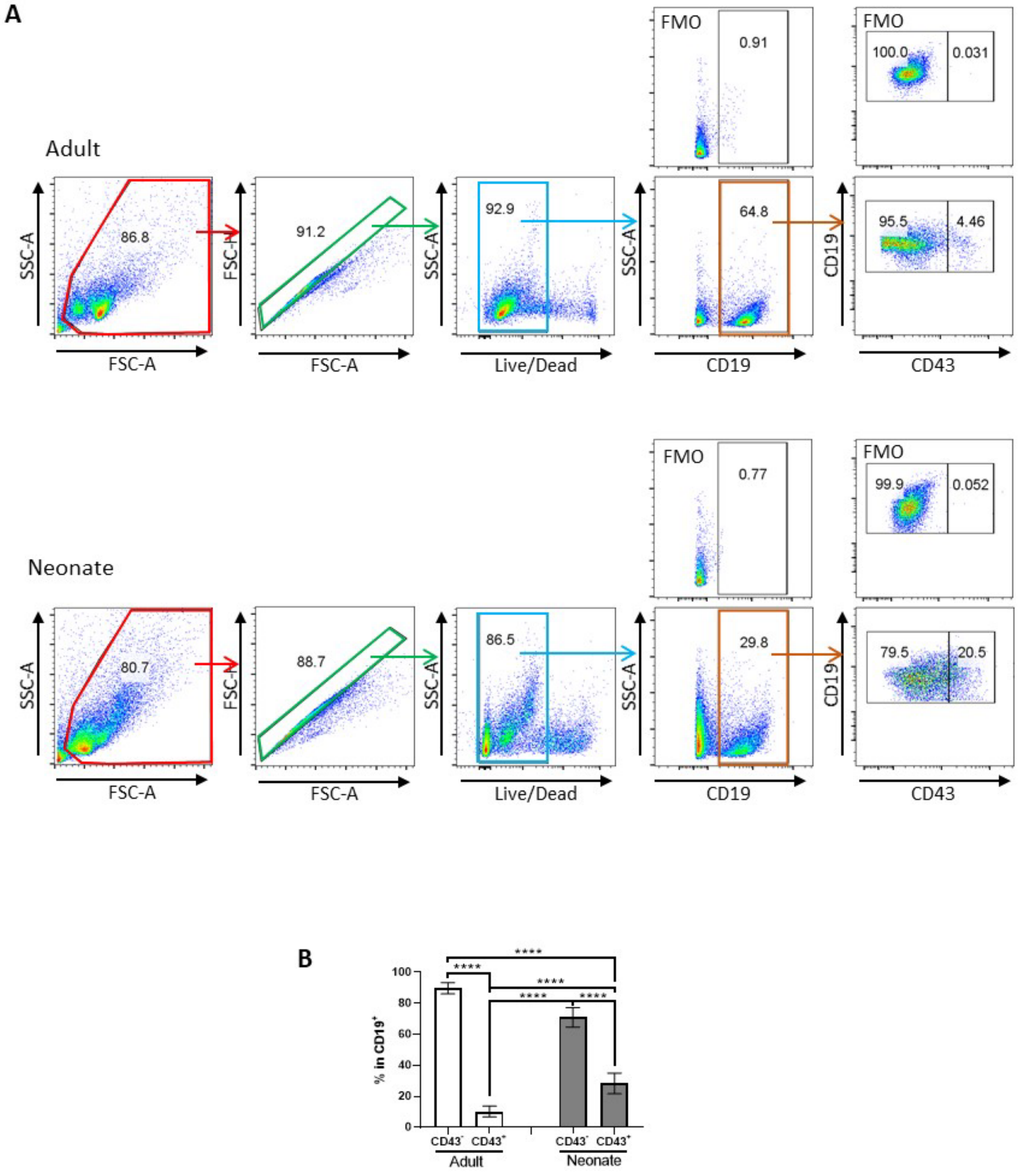


**Supplementary Figure S2. Staining of splenic B cells subsets for IL-10 expression. (A)** Flow cytometry gating strategy for identification of splenic CD19$^{+}$CD43$^{+}$ B cells and CD19$^{+}$CD43$^{-}$ B cells. **(B)** Splenocytes were analyzed for CD19$^{+}$CD43$^{+}$ B cells and CD19$^{+}$CD43$^{-}$ B cells.in adult and neonatal spleen. Data are shown as the mean ± s.d. of three independent experiments. P values were calculated using one-way ANOVA with a Dunnett's multiple comparisons test (*****P*<0.0001).

**Figure S3**

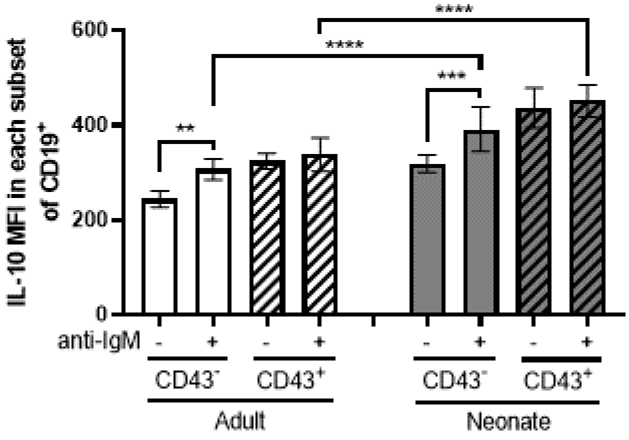


**Supplementary Figure S3. BCR-induced IL-10 production in adult and neonatal splenic B cell subsets.** IL-10 MFI of each subset are shown ($n$ = 3). Data are shown as the mean ± s.d. of three independent experiments. $P$ values were calculated using one-way ANOVA with a Dunnett's multiple comparisons test (**$P$<0.01, ***$P$<0.001 and ****$P$<0.0001).

**Figure S4**

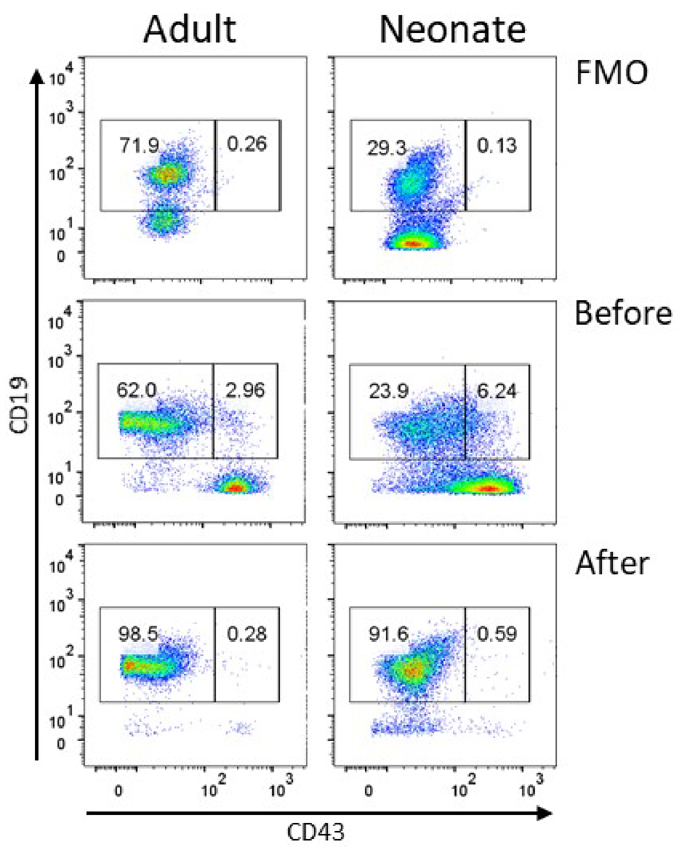


**Supplementary Figure S4. Isolation of splenic CD19$^{+}$CD43$^{-}$ B cells.** Splenic CD19$^{+}$CD43$^{-}$ B cells were isolated using B Cell Isolation Kit (130-090-862, Miltenyi Biotec) which contains anti-CD43 antibody as well as anti-CD4 and anti-Ter119 antibody.

**Figure S5**

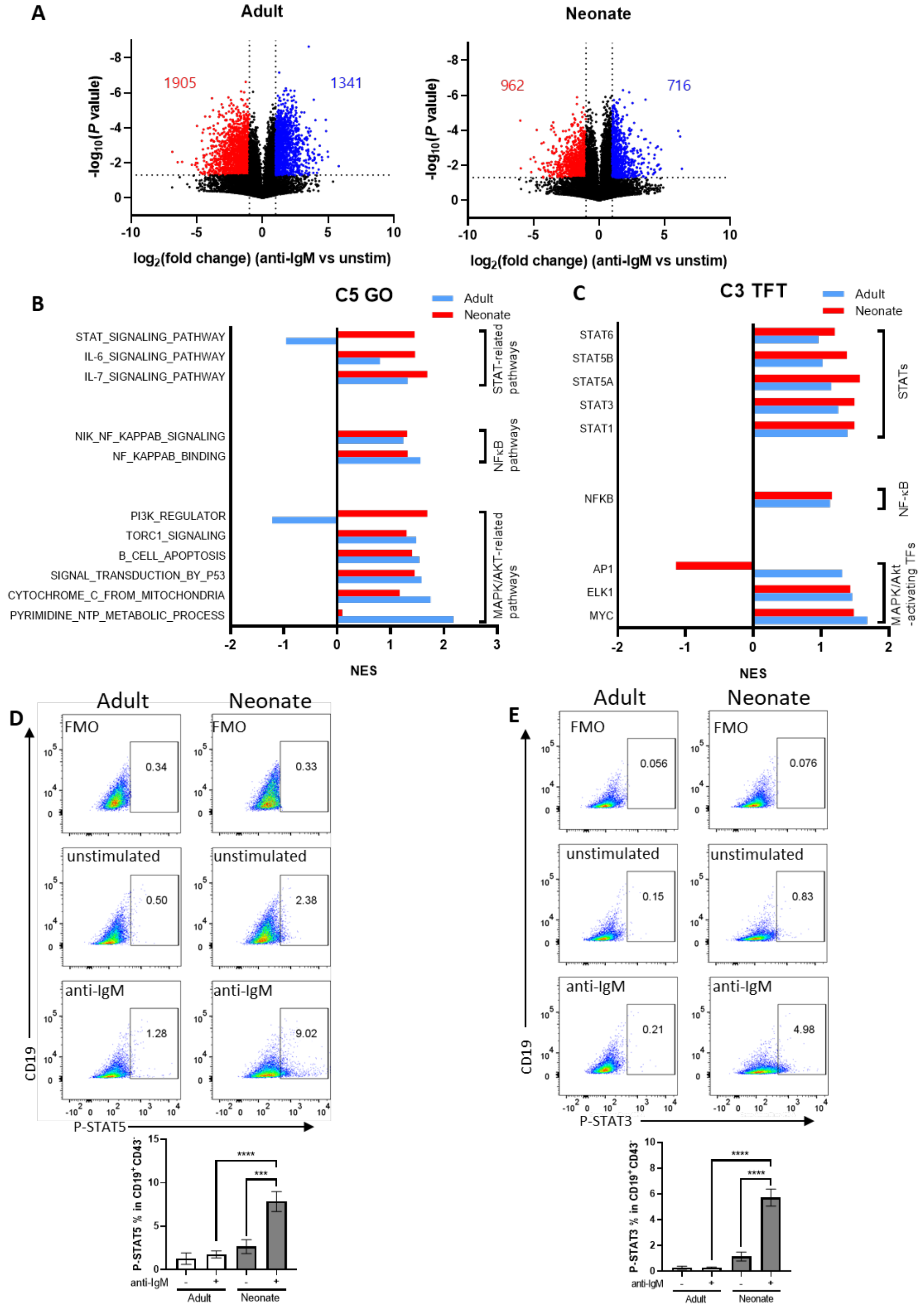
A
Adult
Neonate
1905
1341
962
716
-log10(P value)
log2(fold change) (anti-IgM vs unstim)
B
C5 GO
Adult
Neonate
STAT_SIGNALING_PATHWAY
IL-6_SIGNALING_PATHWAY
IL-7_SIGNALING_PATHWAY
NIK_NF_KAPPAB_SIGNALING
NF_KAPPAB_BINDING
PI3K_REGULATOR
TORC1_SIGNALING
B_CELL_APOPTOSIS
SIGNAL_TRANSDUCTION_BY_P53
CYTOCHROME_C_FROM_MITOCHONDRIA
PYRIMIDINE_NTP_METABOLIC_PROCESS
STAT-related pathways
NFκB pathways
MAPK/AKT-related pathways
NES
C
C3 TFT
STAT6
STAT5B
STAT5A
STAT3
STAT1
NFKB
AP1
ELK1
MYC
STATs
NF-κB
MAPK/Akt -activating TFs
NES
D
Adult
Neonate
FMO
0.34
0.33
unstimulated
0.50
2.38
anti-IgM
1.28
9.02
CD19
P-STAT5
P-STAT5 % in CD19+CD43-
****
***
anti-IgM
Adult
Neonate
E
Adult
Neonate
FMO
0.056
0.076
unstimulated
0.15
0.83
anti-IgM
0.21
4.98
CD19
P-STAT3
P-STAT3 % in CD19+CD43-
****
****
anti-IgM
Adult
Neonate

**Supplementary Figure S5. STAT3 and STAT5 are uniquely activated in neonatal $CD19^+CD43^-$ B cells following BCR cross-linking.** Isolated $CD19^+CD43^-$ cells were stimulated with 10 µg/mL $F(ab')_2$ fragments of anti-IgM antibodies for 7 h, and then total RNA was isolated for regular RNA sequencing. **(A)** Volcano plots of the differential gene expression levels in adult and neonatal B cells following BCR cross-linking are shown. **(B)** Pathway analysis of signaling pathways upregulated following BCR activation are shown. GSEA was performed using the C5 gene ontology (GO) gene sets in MSigDB. NES for the changes in the same pathways are compared between Adult and Neonatal cells. **(C)** GSEA analysis was performed using the hallmark gene sets in C3 transcription factor targets (TFT). NES for each TF are compared between Adult and Neonatal cells. **(D and E)** Flow cytometry analysis of STAT5 phosphorylation **(D)** and STAT3 phosphorylation **(E)** was done on stimulated $CD19^+CD43^-$ B cells. Isolated splenic $CD19^+CD43^-$ B cells were unstimulated (unstim) or stimulated with 10 µg/mL $F(ab')_2$ fragments of anti-IgM antibodies for 15 min **(D)** or 4 h **(E)** and phospho-STAT5 and phosphor-STAT3 were measured ($n$ = 3). Data are shown as the mean ± s.d. of three independent experiments. ***$P$<0.001 and ****$P$<0.0001. $P$ values were calculated using one-way ANOVA with a Dunnett's multiple comparisons test.

**Figure S6**

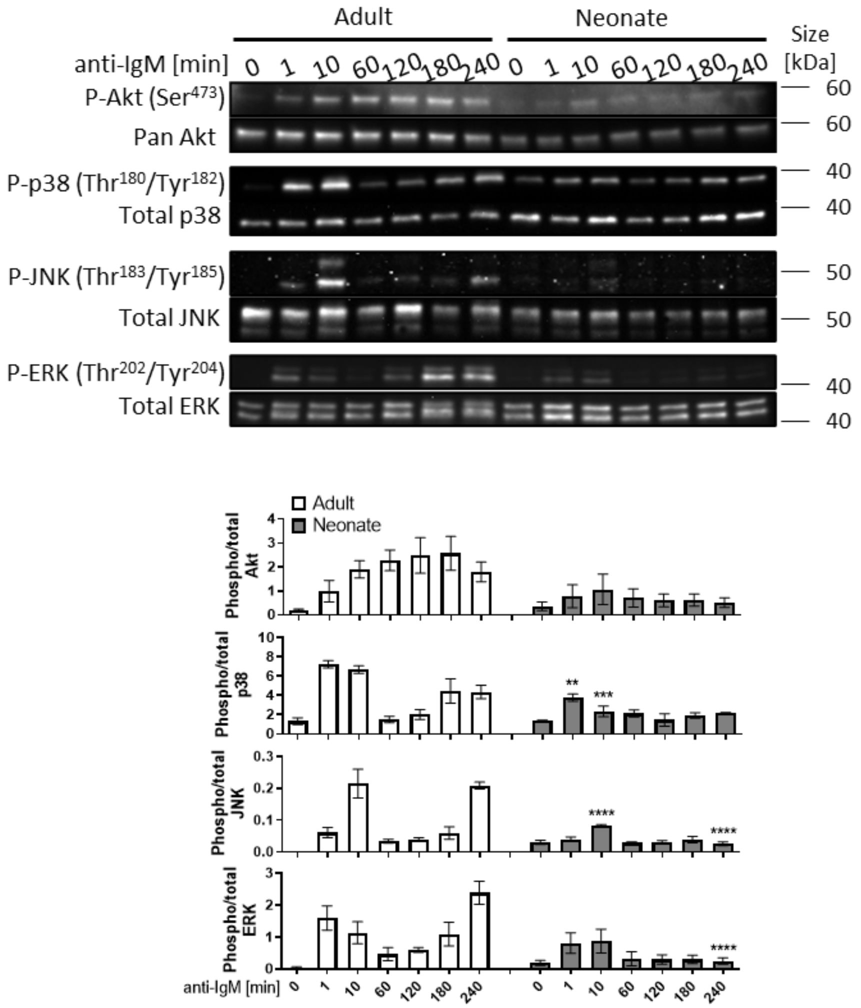


**Supplementary Figure S6. Akt, p38, JNK, and ERK phosphorylations in adult and neonatal splenic CD19$^+$CD43$^-$ B cells following BCR cross-linking.** Splenic CD19$^+$CD43$^-$ cells isolated from adult and neonatal mice were stimulated with 10 μg/mL F(ab')$_2$ fragments of anti-IgM antibodies for the indicated duration, and whole cell extracts were collected for immunoblot analysis of Akt, p38, JNK and ERK. Data are shown as the mean ± s.d. of three independent experiments. *P* values were calculated using one-way ANOVA with a Dunnett's multiple comparisons test (***P*<0.01, ****P*<0.001 and *****P*<0.0001).

### Figure S7

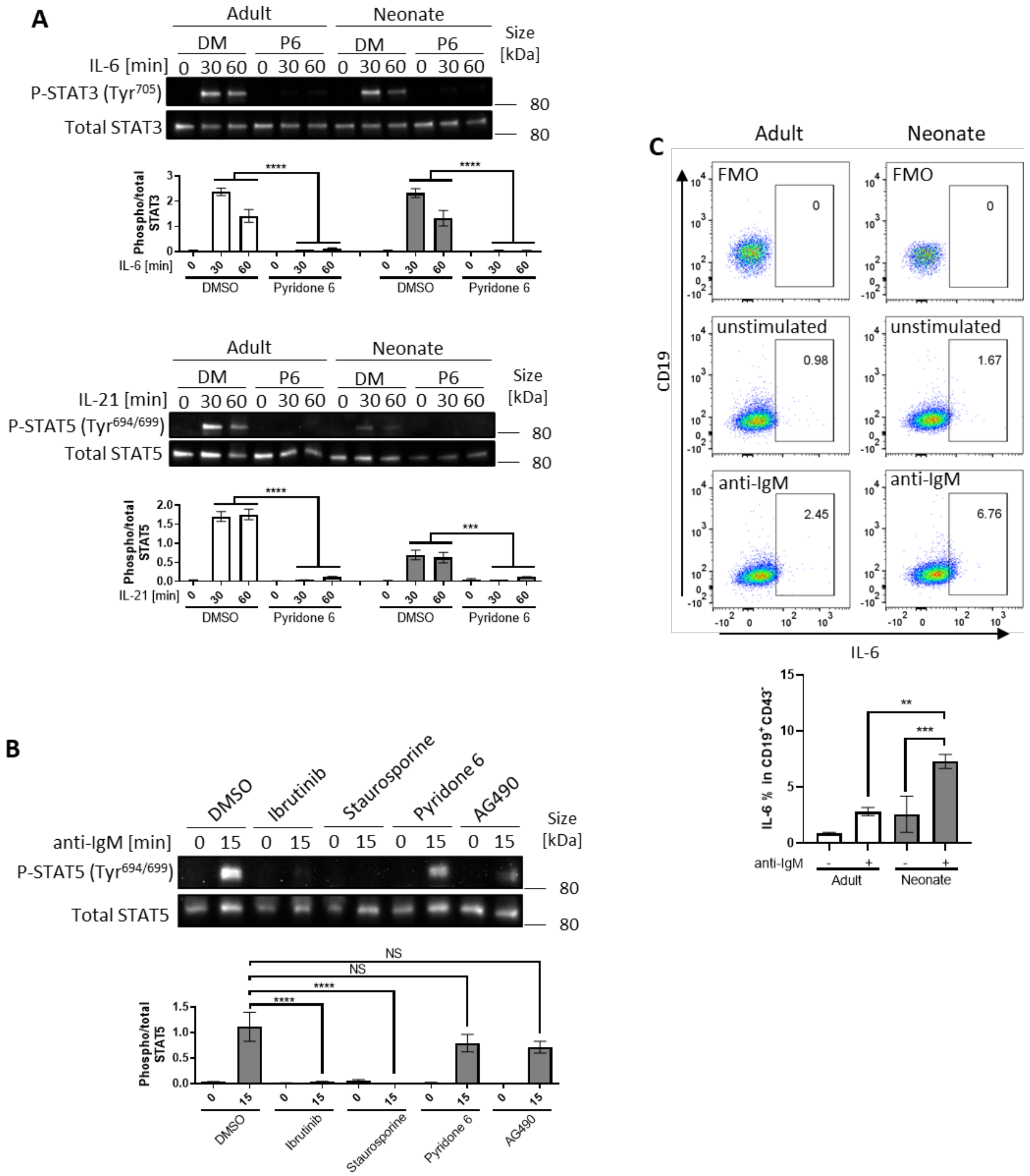
A
Adult
Neonate
DM
P6
IL-6 [min]
0 30 60
P-STAT3 (Tyr705)
Total STAT3
Size [kDa]
80
Phospho/total STAT3
DMSO
Pyridone 6
IL-21 [min]
P-STAT5 (Tyr694/699)
Total STAT5
Phospho/total STAT5
B
DMSO
Ibrutinib
Staurosporine
Pyridone 6
AG490
anti-IgM [min]
0 15
NS
C
FMO
unstimulated
anti-IgM
0
0.98
1.67
2.45
6.76
CD19
IL-6
IL-6 % in CD19+CD43-
anti-IgM

**Supplementary Figure S7. The effect of inhibitors on STAT3 and STAT5 phosphorylations in splenic CD19$^+$CD43$^-$ cells. (A)** Isolated splenic CD19$^+$CD43$^-$ cells were pre-treated with DMSO (DM) or 1 μM Pyridone 6 (P6) for 1 h prior to stimulation with 20 ng/mL recombinant IL-6 or 20 ng/mL recombinant IL-21 for the indicated duration and phosphorylation of STAT3 and STAT5 were investigated in Western blot analysis. Data are shown as the mean ± s.d. of three independent experiments. *P* values were calculated using one-way ANOVA with a Dunnett's multiple comparisons test (***$P$<0.001 and ****$P$<0.0001). **(B)** Splenic CD19$^+$CD43$^-$ cells isolated from neonatal mice were pre-treated with DMSO, BTK inhibitor Ibrutinib, PKC inhibitor Staurosporine, pan-JAK inhibitor Pyridone 6, or JAK2 inhibitor AG90 for 1 h prior to stimulation with 10 μg/mL F(ab')$_2$ fragments of anti-IgM antibodies for the indicated duration. Phosphorylation of STAT5 was determined in Western blot analysis. Data are shown as the mean ± s.d. of three independent experiments. *P* values were calculated using one-way ANOVA with a Dunnett's multiple comparisons test (NS: no significant difference, and ****$P$<0.0001). **(C)** Isolated splenic CD19$^+$CD43$^-$ B cells were unstimulated or stimulated with 10 μg/mL F(ab')$_2$ fragments of anti-IgM antibodies for 17 h and intracellular IL-6 were measured in CD19$^+$CD43$^-$ B cells ($n$ = 3). Data are shown as the mean ± s.d. of three independent experiments. *P* values were calculated using one-way ANOVA with a Dunnett's multiple comparisons test (**$P$<0.01, and ***$P$<0.001).

**Figure S8**

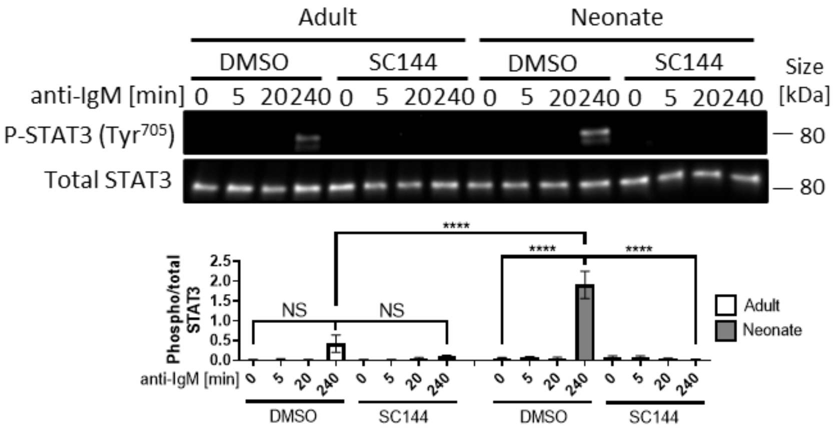


**Supplementary Figure S8. Inhibition of gp130 blocks STAT3 activation in BCR stimulated neonatal cells.** Isolated splenic $CD19^{+}CD43^{-}$ cells were pre-treated with DMSO, 2 μM gp130 inhibitor SC144 for 1 h prior to stimulation with 10 μg/mL $F(ab')_2$ fragments of anti-IgM antibodies for the indicated duration. Phosphorylation of STAT3 was determined in Western blot analysis. Data are shown as the mean ± s.d. of three independent experiments. *P* values were calculated using one-way ANOVA with a Dunnett's multiple comparisons test (NS: no significant difference, *****P*<0.0001).

**Figure S9**

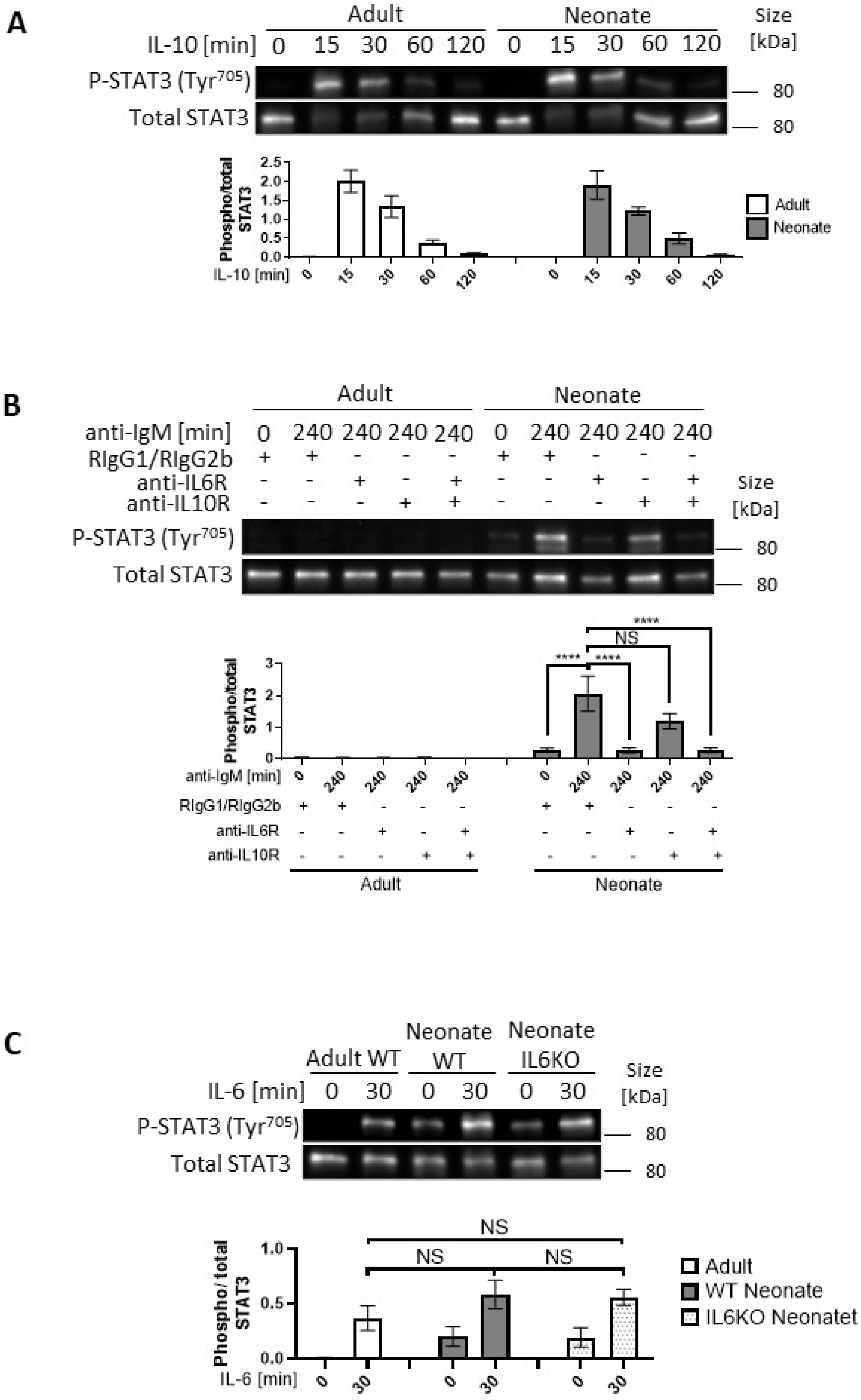
A
Adult
Neonate
IL-10 [min] 0 15 30 60 120 0 15 30 60 120
Size [kDa]
P-STAT3 (Tyr705)
Total STAT3
80
80
Phospho/total STAT3
Adult
Neonate
B
Adult
Neonate
anti-IgM [min] 0 240 240 240 240 0 240 240 240 240
RIgG1/RIgG2b
anti-IL6R
anti-IL10R
Size [kDa]
P-STAT3 (Tyr705)
Total STAT3
80
80
Phospho/total STAT3
****
NS
C
Adult WT
Neonate WT
Neonate IL6KO
IL-6 [min] 0 30 0 30 0 30
Size [kDa]
P-STAT3 (Tyr705)
Total STAT3
80
80
Phospho/ total STAT3
NS
Adult
WT Neonate
IL6KO Neonatet

**Supplementary Figure S9. Autocrine IL-10 has a minor role in STAT3 activation following BCR cross-linking. (A)** Isolated splenic $CD19^{+}CD43^{-}$ cells were stimulated with 10 nM recombinant IL-10 for the indicated duration and STAT3 phosphorylation was assessed in Western blot analysis. Data are shown as the mean ± s.d. of three independent experiments. *P* values were calculated using one-way ANOVA with a Dunnett's multiple comparisons test. **(B)** Isolated $CD19^{+}CD43^{-}$ B cells were pre-treated with 10 μg/mL isotype control antibodies (Rat IgG1 and Rat IgG2b), anti-IL-6R antibody and anti-IL-10R antibody for 1 h prior to stimulation with 10 μg/mL $F(ab')_2$ fragments of anti-IgM antibodies for the indicated duration and STAT3 phosphorylation was assessed in Western blot analysis. Data are shown as the mean ± s.d. of three independent experiments. *P* values were calculated using one-way ANOVA with a Dunnett's multiple comparisons test (NS: no significant difference, and ****$P$<0.0001). **(C)** Isolated $CD19^{+}CD43^{-}$ B cells from wild-type (WT) adult, WT neonate and IL-6-deficient (IL6KO) neonates were stimulated with 20 ng/mL recombinant IL-6 for the indicated duration. STAT3 phosphorylation was determined in Western blot analysis. Data are shown as the mean ± s.d. of three independent experiments. *P* values were calculated using one-way ANOVA with a Dunnett's multiple comparisons test (NS: no significant difference).

**Figure S10**

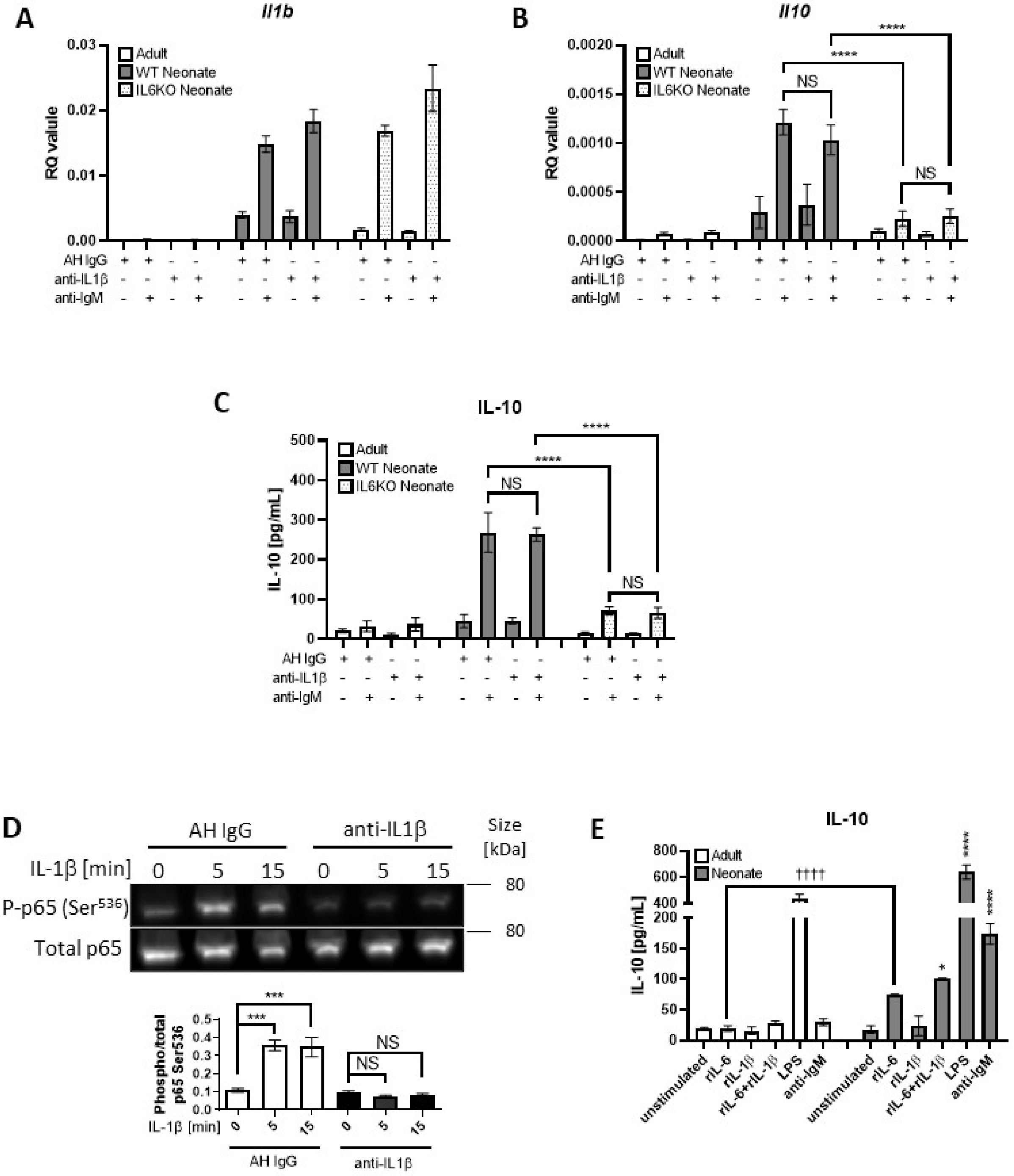
A
Il1b
RQ value
Adult
WT Neonate
IL6KO Neonate
AH IgG
anti-IL1β
anti-IgM
B
Il10
NS
****
C
IL-10
IL-10 [pg/mL]
D
AH IgG
anti-IL1β
Size [kDa]
IL-1β [min]
P-p65 (Ser536)
Total p65
80
Phospho/total p65 Ser536
***
NS
E
IL-10
Neonate
††††
unstimulated
rIL-6
rIL-1β
rIL-6+rIL-1β
LPS
anti-IgM
*

**Supplementary Figure S10. Autocrine/paracrine IL-1β is not involved in neonatal BCR-induced IL-10 production. (A-C)** Splenic $CD19^+CD43^-$ cells isolated from wild-type (WT) adult, WT neonate, and IL-6-deficient (IL6KO) neonate were treated with 10 μg/mL isotype control antibody (American Hamster IgG) or anti-IL1β antibody for 1 h prior to stimulation with 10 μg/mL $F(ab')_2$ fragments of anti-IgM antibodies for 18 h. *Il1b* **(A)** ($n$ = 3) and *Il10* **(B)** ($n$ = 3) mRNA expression were determined by RT-qPCR and the secreted IL-10 were determined by ELISA **(C)** ($n$ = 3). Data are shown as the mean ± s.d. $P$ values were calculated using one-way ANOVA with a Dunnett's multiple comparisons test (****$P$<0.0001). **(D)** Splenic $CD19^+CD43^-$ cells isolated from adult were treated with 10 μg/mL isotype control antibody (Armenian Hamster IgG) or anti-IL1β antibody for 1 h prior to stimulation with 20 ng/mL recombinant IL-1β for the indicated duration. Data are shown as the mean ± s.d. of three independent experiments. $P$ values were calculated using one-way ANOVA with a Dunnett's multiple comparisons test (***$P$<0.001). **(E)** Splenic $CD19^+CD43^-$ cells isolated from adult and neonatal mice were stimulated with 20 ng/mL recombinant IL-6, 20 ng/mL recombinant IL-1β, 10 μg/mL LPS, or 10 μg/mL $F(ab')_2$ fragments of anti-IgM antibodies for 8 h. Culture supernatant IL-10 levels were determined by ELISA ($n$ = 3). Data are shown as the mean ± s.d. $P$ values versus adult counterparts were calculated using one-way ANOVA with a Dunnett's multiple comparisons test (*$P$<0.05 and ****$P$<0.0001) and two-tailed unpaired $t$-tests (††††$P$<0.0001).

**Figure S11**

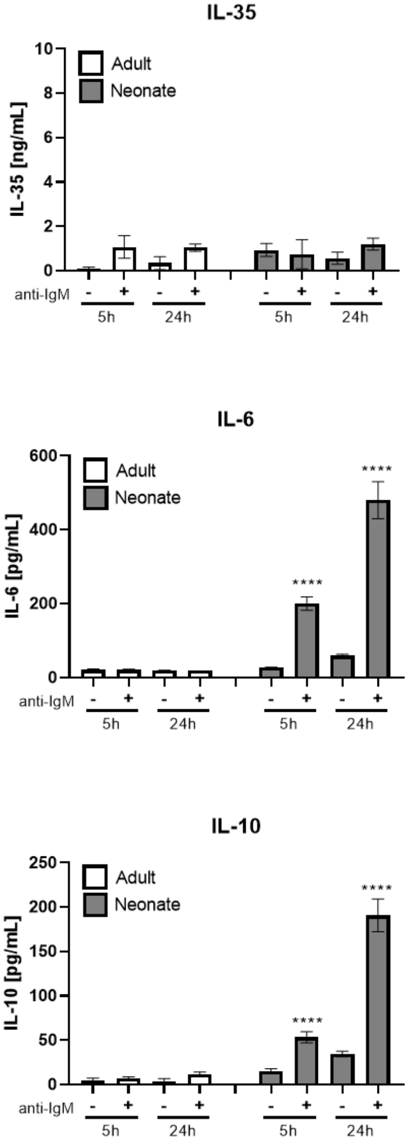


**Supplementary Figure S11 Autocrine/paracrine IL-35 is not involved in neonatal BCR-induced IL-10 production.** Splenic $CD19^{+}CD43^{-}$ cells isolated from adult and neonatal mice were stimulated with 10 μg/mL $F(ab')_2$ fragments of anti-IgM antibodies for 5 h and 24 h. Culture supernatant IL-35, IL-6 and IL-10 levels were determined by ELISA ($n$ = 3). Data are shown as the mean ± s.d. of two independent experiments. $P$ values versus adult counterparts were calculated using one-way ANOVA with a Dunnett's multiple comparisons test (****$P$<0.0001).

**Figure S12**

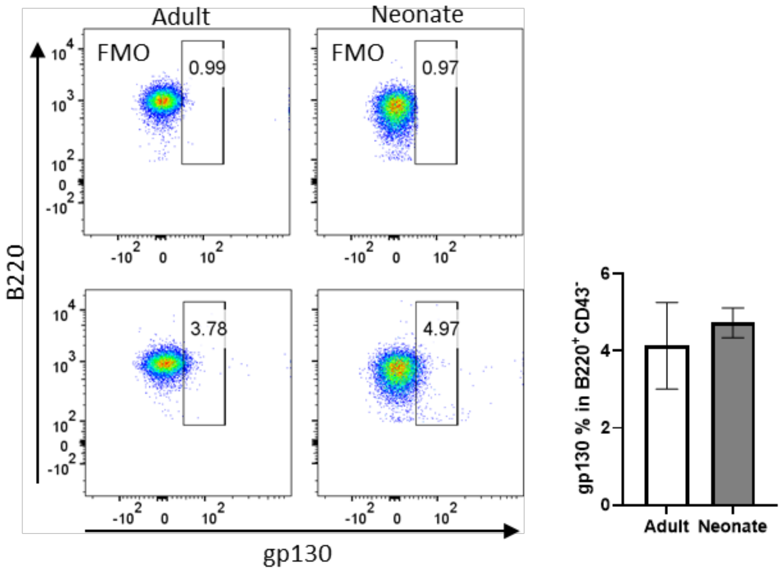


**Supplementary Figure S12. STAT3 is activated by IL-6 in an autocrine fashion.** Isolated CD19$^{+}$CD43$^{-}$ B cells were analyzed for surface levels of gp130 ($n$ = 3). Data are shown as the mean ± s.d. of three independent experiments. $P$ values versus adult counterparts were calculated using two-tailed unpaired $t$-tests (*$P$<0.05).

**Figure S13**

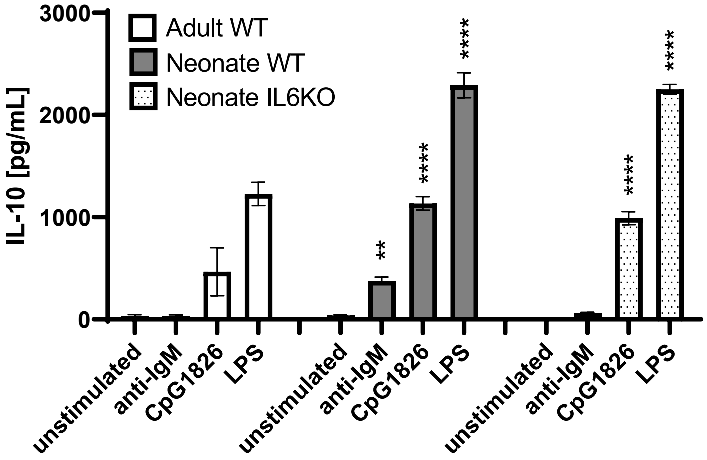


**Supplementary Figure S13. TLR-induced IL-10 production is not dependent on IL-6.** Splenic $CD19^{+}CD43^{-}$ cells isolated from wild-type (WT) adult, WT neonate, and IL-6-deficient neonate (IL6KO) were stimulated with 10 μg/mL $F(ab')_2$ fragments of anti-IgM antibodies, 10 μg/mL CpG 1826, or 10 μg/mL lipopolysaccharide (LPS) for 18 h. Culture supernatant IL-10 levels were determined by ELISA ($n$ = 3). Data are shown as the mean ± s.d. of two independent experiments. $P$ values versus adult counterparts were calculated using one-way ANOVA with a Dunnett's multiple comparisons test (**$P$<0.01 and ****$P$<0.0001).

**Figure S14**

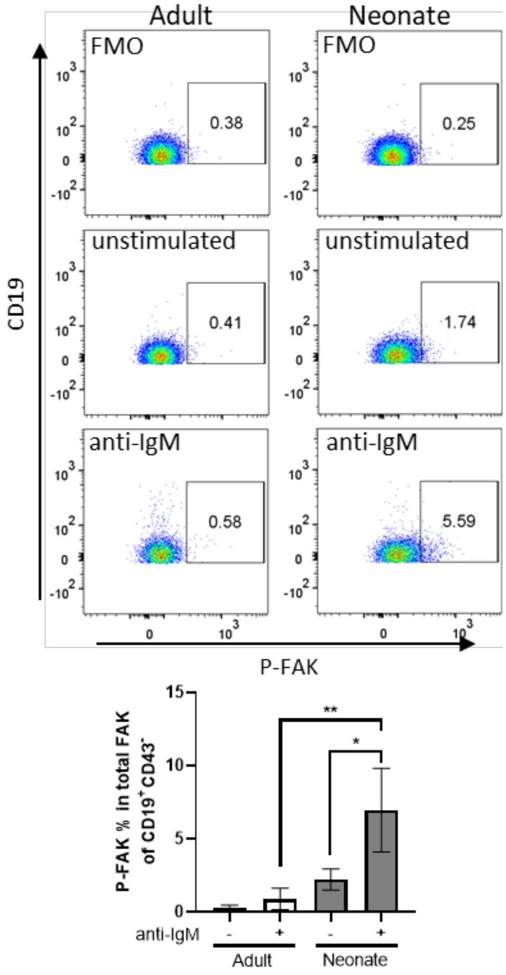


**Supplementary Figure S14. Neonatal BCRs activate FAK.** Isolated splenic $CD19^{+}CD43^{-}$ B cells were unstimulated or stimulated with 10 μg/mL F(ab')$_2$ fragments of anti-IgM antibodies for 30 sec and intracellular phospho-FAK was measured in $CD19^{+}CD43^{-}$ B cells (*n* = 3). Data are shown as the mean ± s.d. of three independent experiments. *P* values were calculated using one-way ANOVA with a Dunnett's multiple comparisons test (**P*<0.05 and ***P*<0.01).

### Figure S15

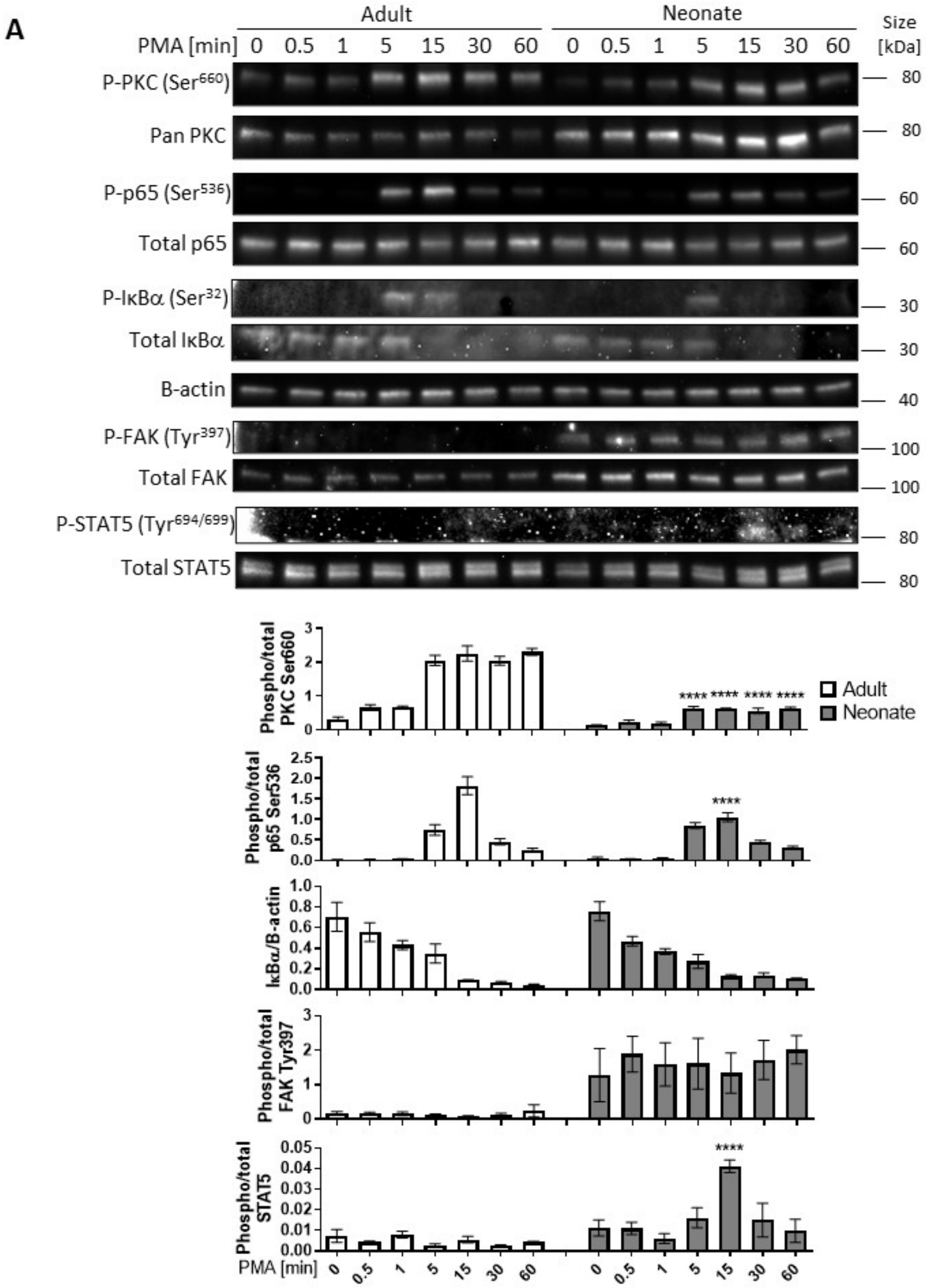
A
Adult
Neonate
PMA [min] 0 0.5 1 5 15 30 60 0 0.5 1 5 15 30 60
Size [kDa]
P-PKC (Ser660)
Pan PKC
P-p65 (Ser536)
Total p65
P-IκBα (Ser32)
Total IκBα
B-actin
P-FAK (Tyr397)
Total FAK
P-STAT5 (Tyr694/699)
Total STAT5
Phospho/total PKC Ser660
Phospho/total p65 Ser536
IκBα/B-actin
Phospho/total FAK Tyr397
Phospho/total STAT5
Adult
Neonate
****

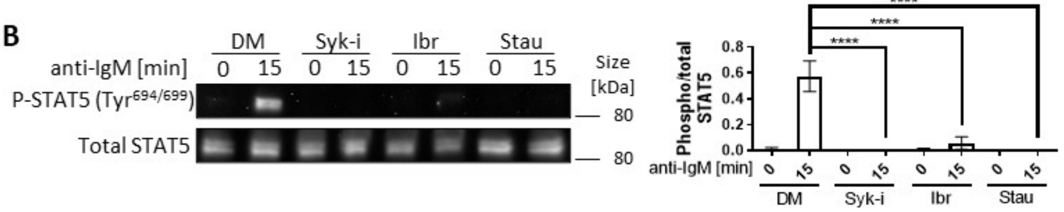
B
DM
Syk-i
Ibr
Stau
anti-IgM [min] 0 15 0 15 0 15 0 15
Size [kDa]
P-STAT5 (Tyr694/699)
Total STAT5
Phospho/total STAT5
****

**Supplementary Figure S15. BCR-induced STAT5 activation is dependent on Syk, Btk, and PKC. (A)** Isolated $CD19^{+}CD43^{-}$ B cells were stimulated with 10 µg/mL PMA for the indicated duration and phosphorylation of PKC, p65, IκBα, FAK and STAT5 were determined in Western blot analysis. Data are shown as the mean ± s.d. of three independent experiments. *P* values were calculated using one-way ANOVA with a Dunnett's multiple comparisons test (**** $P<0.0001$). **(B)** Isolated $CD19^{+}CD43^{-}$ cells from neonatal spleen were pre-treated with DMSO, 10 µM Syk inhibitor (Syk-i), 5 µM Btk inhibitor Ibrutinib (Ibr), or 20 nM Staurosporine for 1 h and then stimulated with 10 µg/mL $F(ab')_2$ fragments of anti-IgM antibodies for the indicated duration and phosphorylation of STAT5 was determined in Western blot analysis. Data are shown as the mean ± s.d. of three independent experiments. *P* values were calculated using one-way ANOVA with a Dunnett's multiple comparisons test (**** $P<0.0001$).

**Figure S16**

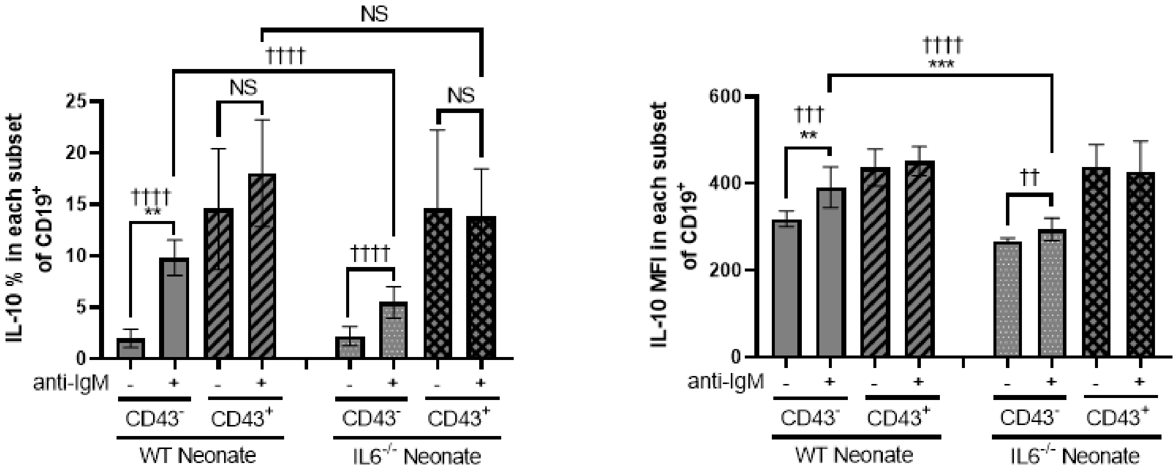


**Supplementary Figure S16. IL-6 is not responsible for IL-10 production by CD43-expressing B1 subset in neonatal mouse.** Isolated CD19$^{+}$ cells were incubated in the absence or presence of 10 μg/mL F(ab')$_2$ fragments of anti-IgM for 20 h (*n* = 3). Data are shown as the mean ± s.d. of three independent experiments. P values were calculated using one-way ANOVA with a Dunnett's multiple comparisons test (***P*<0.01, NS: no significance) and two-tailed unpaired *t*-tests (††*P*<0.01, †††*P*<0.001 and ††††*P*<0.0001).

**Figure S17**

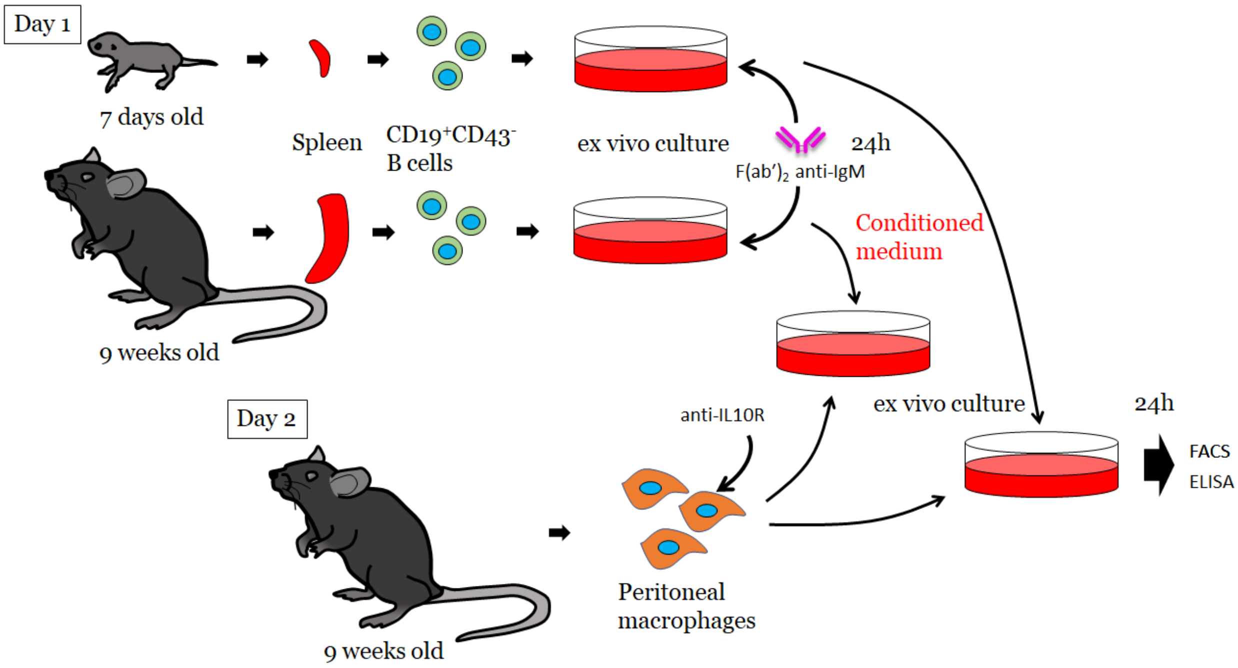


**Supplementary Figure S17. A schematic description of experimental design to test the suppressive effect of IL-10 secreted from neonatal $CD19^{+}CD43^{-}$ B cells.** Isolated splenic $CD19^{+}CD43^{-}$ B cells from adult and neonatal mice were plated at 3.0 million cells per mL and stimulated with 10 μg/mL $F(ab')_2$ fragments of anti-IgM antibodies for 24 h. Conditioned medium were collected and filtered through 0.2 μm membrane. Peritoneal macrophages isolated from adult mice were treated with 10 μg/mL anti-IL10R (1B1.3A) or isotype control antibodies (Rat IgG1) for 1h prior to culture in conditioned medium. After 24h incubation, supernatants were collected for the assessments.

**Figure S18**

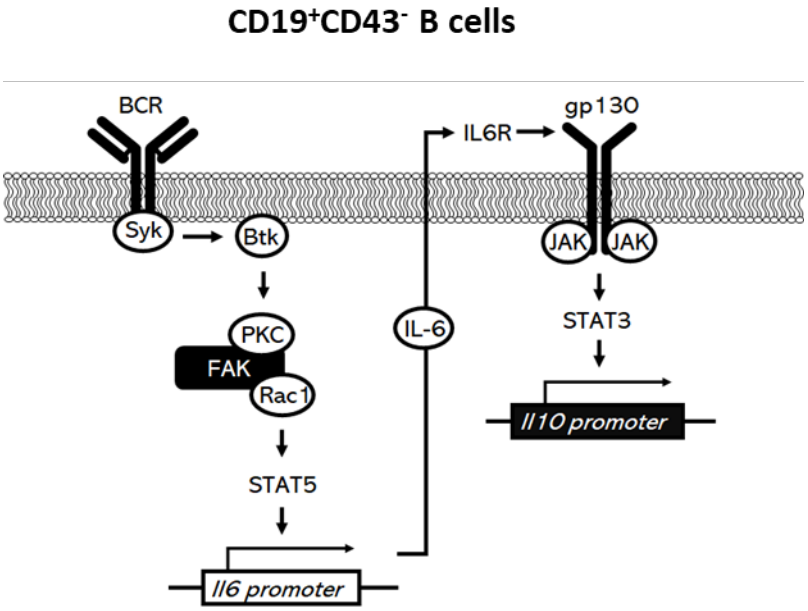


**Supplementary Figure S18. Schematic representation of IL-10 production in BCR stimulated neonatal $CD19^+CD43^-$ cells.** Recognition of antigen by BCR induces IL-6 secretion from $CD19^+CD43^-$ B cells through a pathway involving Syk, Btk, PKC, FAK, Rac1 and STAT5. The secreted IL-6 binds to its receptors IL-6Rα and gp130 on $CD19^+CD43^-$ B cells to stimulate the production of IL-10 in a JAK and STAT3 dependent manner.
